# Multi-model inference of non-random mating from an information theoretic approach

**DOI:** 10.1101/305730

**Authors:** Antonio Carvajal-Rodríguez

## Abstract

Non-random mating has a significant impact on the evolution of organisms. Here, I developed a modelling framework for discrete traits (with any number of phenotypes) to explore different models connecting the non-random mating causes (mate competition and/or mate choice) and their consequences (sexual selection and/or assortative mating).

I derived the formulas for the maximum likelihood estimates of each model and used information criteria to perform multi-model inference. Simulation results showed a good performance of both model selection and parameter estimation. The methodology was applied to ecotypes data of the marine gastropod *Littorina saxatilis* from Galicia (Spain), to show that the mating pattern is better described by models with two parameters that involve both mate choice and competition, generating positive assortative mating plus female sexual selection. As far as I know, this is the first standardized methodology for model selection and multi-model inference of mating parameters for discrete traits. The advantages of this framework include the ability of setting up models from which the parameters connect causes, as mate competition and mate choice, with their outcome in the form of data patterns of sexual selection and assortative mating. For some models, the parameters may have a double effect i.e. they produce sexual selection and assortative mating, while for others there are separated parameters for one kind of pattern or another. From an empirical point of view, it is much easier to study patterns than processes and, for this reason, the causal mechanisms of sexual selection are not so well known as the patterns they produce. The goal of the present work is to propose a new tool that helps to distinguish among different alternative processes behind the observed mating pattern.

The full methodology was implemented in a software called InfoMating (available at http://acraaj.webs6.uvigo.es/InfoMating/Infomating.htm).

## 1. Introduction

The concept of sexual selection is a key piece of modern evolutionary theory as it explains a great range of evolutionary patterns and diversity. Darwin (1871, 1974) originally defined sexual selection as competition between individuals of one sex to achieve matings with the other sex. Yet Darwin distinguished two general biological mechanisms of sexual selection: mate competition and mate choice (see Ng et al. 2019 and references therein). However, the concept of sexual selection has been controversial since its very beginning (reviewed in Andersson 1994; Prum 2012; Parker 2014; Parker and Pizzari 2015) and there is still disagreement on its actual definition (Fitze and Galliard 2011), and even, its role as a key component of modern evolutionary biology has being challenged (Roughgarden et al. 2006; but see Shuker 2010; Parker and Pizzari 2015).

It seems that some of the disagreements and misunderstandings about sexual selection and related concepts, come from the distinct emphases that scientific fields (e.g. population genetics, speciation theory, behavioral ecology and sociology) put on the various aspects of the sexual selection theory (evolutionary, behavioral or social role). Moreover, sexual selection is described sometimes as a process and sometimes as a pattern.

The distinction between pattern and process may be obscured because of the same biological concept can be meaningfully defined as both a process and a pattern (Armstrong 1977; Mahler et al. 2017). Consider for example, the classical definition of sexual selection, as arising from variation in reproductive success due to competition for access to mates (Andersson 1994; Shuker 2010). From such definition, sexual selection can be considered as the evolutionary agent (a process) that drives the evolution of some mating-related traits. However, from the same definition, if we put the emphasis on the pattern of evolutionary change that arises from the differences in the reproductive success, then we are viewing sexual selection as a pattern caused by some other biological process (competition).

In this work, I adhere to the definition used in population genetics, where sexual selection is caused by processes of mate competition that may produce intrasexual selection, and/or processes of mate choice that may produce intersexual selection (Lewontin et al. 1968; Endler 1986; Casares et al. 1998; Rolán-Alvarez and Caballero 2000; Ng et al. 2019).

More specifically, the process of mate competition refers in the broad sense to access to matings by courtship, intrasexual aggression and/or competition for limited breeding resources (Andersson 1994; Kokko et al. 2012; Wacker and Amundsen 2014). These processes may generate a pattern of sexual selection (a change in the frequency of the trait under study) in the sex that competes (intrasexual selection Ng et al. 2019).

The process of mate choice occurs whenever the effects of traits expressed in one sex leads to non-random allocation of reproductive investment with members of the opposite sex (Edward 2015). Choice may be mediated by phenotypic (sensorial or behavioural) properties that affect the propensity of individuals to mate with certain phenotypes (Jennions and Petrie 1997). The observed pattern driven by mate choice can be a change in trait frequency in the other sex (intersexual selection) and/or a pattern of trait correlation between mates (assortative mating).

Still, the relationships among these concepts are complex and can be approached from different perspectives (the reader may consult Arnold and Wade 1984; Rolán-Alvarez and Caballero 2000; Edward 2015; Rolan-Alvarez et al. 2015b; Futuyma and Kirkpatrick 2017; Rosenthal 2017; Estévez et al. 2018; Ng et al. 2019; for extended details and alternative definitions).

Summarizing, the evolutionary consequences of mate competition and mate choice are sexual selection and assortative mating. When the traits under study are discrete, the patterns of sexual selection and assortative mating are defined in terms of change in the phenotype frequencies, so that sexual selection corresponds to the observed change in gene or phenotype frequencies in mated individuals with respect to population frequencies (Hartl and Clark 1997; Rolán-Alvarez and Caballero 2000). Similarly, assortative mating corresponds to the observed deviation from random mating within matings (Rolán-Alvarez and Caballero 2000 and references therein).

In a previous work (Carvajal-Rodríguez 2018b), the processes of mate competition and mate choice were modelled for discrete traits by means of the parameters *m*_ij_, that represent the mutual mating propensity between a female of type *i* and a male *j*. Therefore, if A-type females prefer A-type males, this mate choice is modelled as a higher mutual mating propensity between these types as compared with the mutual mating propensity of the A females with other male types (*m*_AA_ > *m*_AB_). On the other hand, if B-type males mate more often than other males whatever the female, this mate competition is modelled by a higher marginal mating propensity of such males (see below).

By modelling the mating process as a differential mutual mating propensity among different types of mating pairs, it is possible to express the mean change in mating phenotypes as the information gained due to non-random mating (Carvajal-Rodríguez 2018b). Describing random mating as the zero information model allows expressing the patterns obtained from mate choice and competition in terms of the information captured in the mutual mating propensity models.

Thus, the mating information-based framework provides a formal approach for developing a set of hypotheses about the causes (mate competition and mate choice) and the patterns they may provoke (sexual selection and assortative mating). In addition, data-based evidence can be used for ranking each hypothesis and perform multi-model-based inference (Link and Barker 2006; Burnham et al. 2011; Aho et al. 2014).

In the following sections I proceed as follows:

1. Given the population frequencies for some discrete trait I define the multinomial saturated mating model in terms of the mutual mating propensity parameters. The maximum-likelihood estimates of these parameters are the pair total indices (PTI) as defined in (Rolán-Alvarez and Caballero 2000). Once the saturated model is defined I obtain the three necessary and sufficient conditions for random mating. Afterwards, by relaxing these conditions it is possible to generate models for which differential marginal mating propensity may produce female or male sexual selection without assortative mating, or on the contrary, models for which some mutual mating propensities represent mate choice that may produce assortative mating and frequency dependent sexual selection. I obtain the maximum likelihood estimates for the parameters of these models.
2. Relying on the previous section, it is possible to generate several mutual mating propensity models and apply information criteria for selecting the best candidate ones and estimating the mating parameter values based on the most supported models. I developed a software called InfoMating to do so.
3. Finally, I demonstrate the methodology by analysing simulated and real data.

## 2. Mutual mating propensity models

Consider a female trait with *k*_1_ different phenotypes and a male trait with *k*_2_ phenotypes, the total number of possible mating phenotypes is *K* = *k*_1_ × *k*_2_. Let a sample have *n*′ matings from which *n*′_ij_ correspond to *i*-type females that mated with *j*-type males. If the probability of the mating *i*×*j* is *q*′_ij_, then the logarithm of the multinomial likelihood function of the sample is

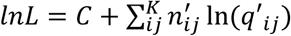

where *C* is the multinomial coefficient which is constant given the sample. As it is well-known, the maximum likelihood estimator of the multinomial probability of the mating *i*×*j* is *n*′_ij_ / *n*′.

### 2.1 Saturated non-random mating model M_sat_

Let the population under study have *n*_1i_ females of type *i* from a total of *n*_1_ females and *n*_2j_ males of type *j* from a total of *n*_2_ males. Therefore, the population frequency of females of type *i* is *p*_1i_ = *n*_1i_ / *n*_1_ and the population frequency of males of type *j* is *p*_2j_ = *n*_2j_ / *n*_2_.

The mating probability between types *i* and *j* can be expressed as *q*′_ij_ = *m*_ij_*q*_ij_ (Carvajal-Rodríguez 2018b) where *q*_ij_ is the product of the female and male population frequencies of each type (*q*_ij_ = *p*_1i_×*p*_2j_) and *m*_ij_ = *m’*_ij_ /*M*, where *m’*_ij_ is the mutual mating propensity, i.e. the expected number of matings given an encounter between females of type *i* and males of type *j*, and *M* is the mean mutual mating propensity *M* = Σ_*i,j*_ *q*_*ij*_*m*′_*ij*_, so that Σ*q*′_ij_ = 1.

Under this multinomial model, the log-likelihood of the sample is

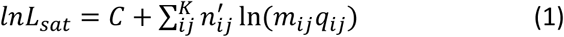

This model is saturated (*M*_sat_) because it has as many parameters as independent mating-class frequencies, *P*_sat_ = *K* −1. The female and male population frequencies, *p*_1_ and *p*_2_, are either known or they need to be estimated in all models. Therefore, for model comparison, the population frequencies can be ignored when counting the number of parameters involved in each model.

The maximum likelihood estimate (MLE) of *m*_ij_ is (*n*′_ij_ / *n*′)/*q*_ij_ = PTI_i,j_ where PTI_i,j_ is the pair total index i.e. the frequency of the observed mating classes divided by their expected frequency under random mating (Rolán-Alvarez and Caballero 2000).

In this work I am interested in the estimation of the mutual mating propensity parameters (from hereafter mutual-propensity parameters) for various competition and mate choice models. From that point of view, it is convenient to express the maximum likelihood estimator in a different way which I call λ-notation.

### 2.2 λ-notation

Consider the non-normalized parameters *m*′_*ij*_ and recall that *m*_ij_ = *m’*_ij_ /*M*. The MLE of *m*′_*ij*_ under *M*_sat_ is simply *M* × PTI_i,j_ i.e. *M* ×(*n*′_ij_ / *n*′) /*q*_ij_ that can be conveniently rearranged as (*n*′_ij_/*q*_ij_)/(*n*′/*M*). Because the mating parameters are normalized, it is possible, without loss of generality, to set just one of the *m*′_*ij*_ to an arbitrary value of 1. Thus, let set *m*′_k1k2_ = 1 and note (details in Appendix A) that in such case *n*′ / *M* = *n*′_k1k2_ / *q*_k1k2_. Therefore, the MLE of the parameters of the saturated model can be expressed as

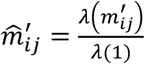

where

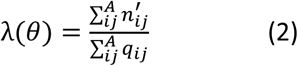

i.e., the function λ of a mating parameter θ is the sum of the counts of all the mating classes in the set *A* = {(*i*_1_, *j*_1_), …} having mutual-propensity θ divided by the sum of their expected frequencies under random mating.

Thus, λ(*m*′_*ij*_) expresses the sum of the observed matings with mutual-propensity *m*′_*ij*_ divided by the product of the population frequencies from each partner type. Similarly, λ(1) corresponds to the sum of the observed matings having unity mating parameter divided by the corresponding products of population frequencies.

As already mentioned, the most parameterized model is the saturated model that has *K*-1 parameters so, when divided by the mean mutual-propensity *M*, the estimates λ(*m*′_*ij*_) / (*M*λ(1)) are the corresponding pair total indices (PTI_ij_).

The model *M*_sat_ is the most complex model that can be fitted to the available data. The principle of parsimony suggests to consider reduced special cases of this saturated model. Next, I computed the ML estimates of different classes of reduced models that require less parameters, beginning by the most reduced one which is the random mating model.

### 2.3 Random mating model M_0_

The random model *M*_0_ corresponds to the simplest, most reduced model, which is nested within all others (it is a particular case of any other model) while it is not possible to derive any simplified version from it. When random mating occurs, the mating probability between types *i* and *j* is *q*′_ij_ = *q*_ij_ = *p*_1i_×*p*_2j_. Under this model, the information would be zero (Carvajal-Rodríguez 2018b). This zero-information model is a particular case of the saturated one when the mutual-propensities are equal for every mating phenotype. The number of independent mating parameters is *P*_0_ = 0.

The log-likelihood of the sample of mating is

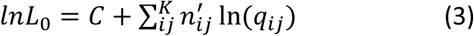

Now, let′s define the marginal propensity *m*_*Fem_i*_ for a female of type *i* as

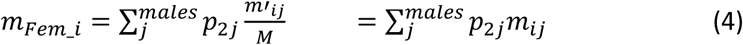

Similarly for a male of type *j*

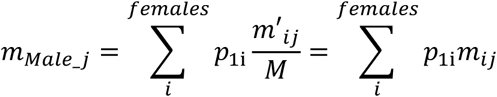

Then, the *M*_0_ model corresponds to *M*_sat_ subjected to the following restrictions:

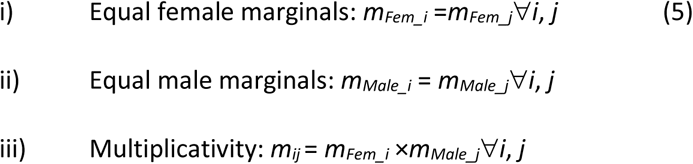

It is useful to express *M*_0_ in terms of these three restrictions because by relaxing some of them it is possible to define different classes of models. For example, a model with equal female marginal propensities and multiplicative mutual-propensities (conditions i and iii hold) but different male marginal propensities (relaxing ii), corresponds to a case with competition among males that may provoke a (intra)sexual selection pattern (see below).

Therefore, by relaxing some of the conditions in (5), it is possible to control the kind of causes that produce the different non-random mating patterns. In fact, there are three general classes of models that can be combined. The two first classes correspond to relaxing the first or second condition and involve mate competition in females or males, provoking female or male (intra)sexual selection respectively. Provided that the third condition is maintained, these models cannot produce an assortative mating pattern (see below). The third class corresponds to relaxing the third condition and involves mate choice, which may provoke just assortative mating, or both assortative mating and sexual selection, the latter depending on the population phenotype frequencies (Fig. 1).

**Fig 1.**
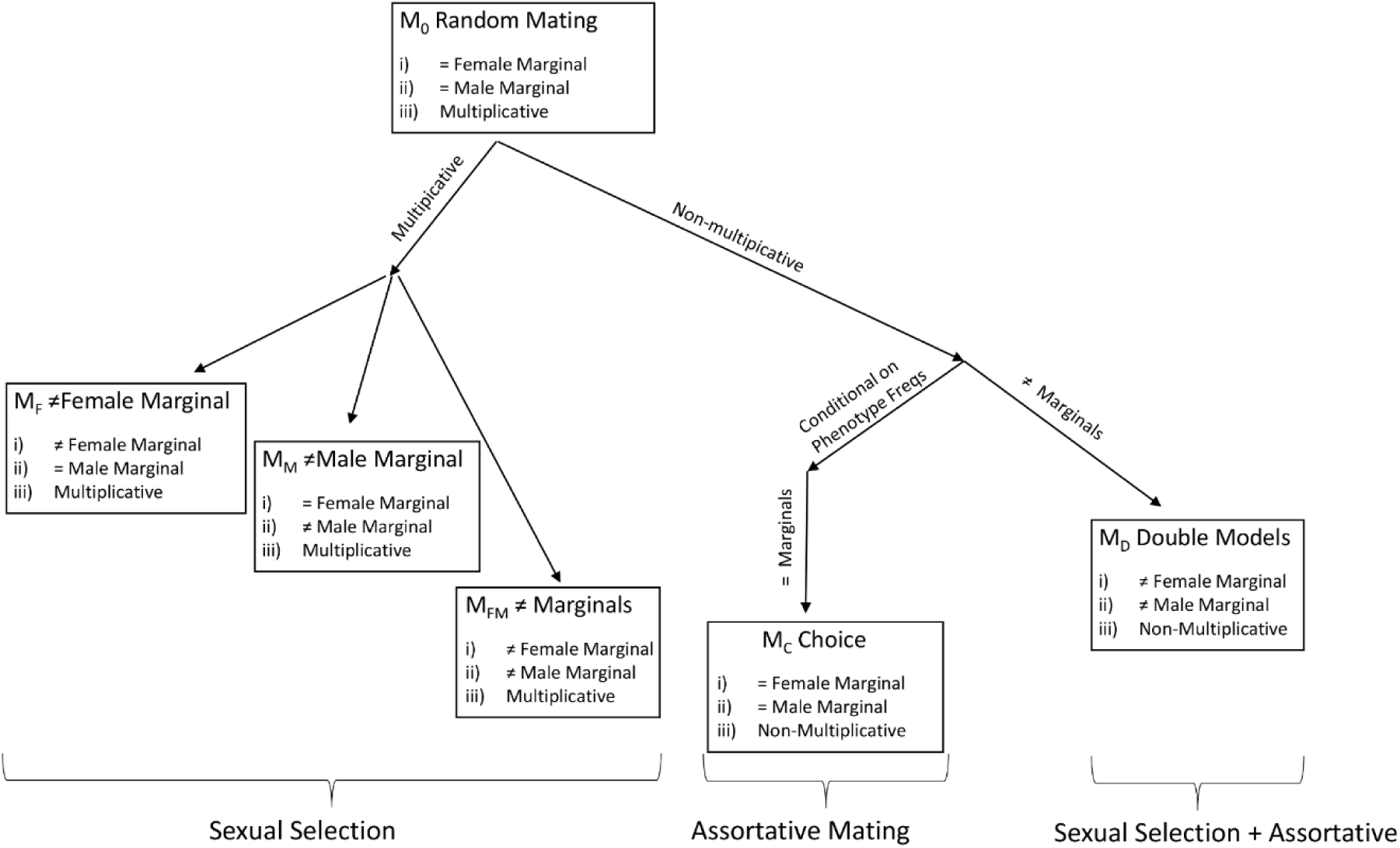
Mating models defined by mate competition or mate choice, and their effect after relaxing some of the conditions imposed to the random mating model *M*_0_.

### 2.4 Mate competition models

These class of models correspond to relaxing the first and/or second conditions in *M*_0_ while maintaining the condition of multiplicativity (5-iii). The maintenance of the third condition implies that the mutual-propensity of a mating pair (*i,j*) is the product of the marginal female (*m*_*Fem*_) and male (*m*_*Male*_) propensities. Under this condition there should be no deviation from random mating when comparing the observed and expected frequencies within matings and the assortative mating pattern should not be observed (Carvajal-Rodríguez 2018b). I distinguished models that generate a sexual selection pattern in just one sex or in both.

#### 2.4.1 Intra-female competition

Relaxing condition (5-*i*) implies that at least one female marginal propensity, say female of type *A*, is different from the rest of female types i.e. *m*_*Fem_A*_ ≠ *m*_*Fem_B*_ with *A* ≠ *B*. On the other side, the marginal propensity of males should be the same which means that there is no intra-male competition, all male types mate at an equal rate.

Therefore, a model with intra-female competition is obtained by defining every mutual-propensity involving a female of type *i*, by an absolute (unnormalized) mating parameter *a*_i_ as follows

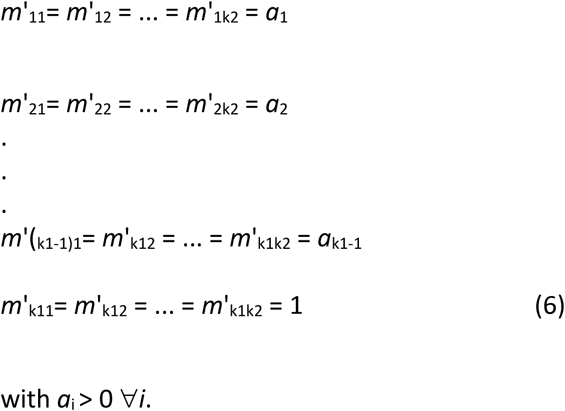

Note that the relationships among the parameters will not be altered when dividing them by *a*_k1_ so that *a*_k1_ = 1. Under this model, there can be as much as *k*_1_-1 free mating parameters.

When computing the female and male marginal propensities (4) it is seen that

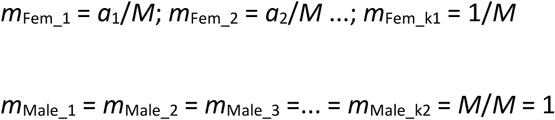

where *M* is the mean mutual-propensity as defined above.

The model (6) has equal male marginal propensity and it is multiplicative. The MLE of the parameters is

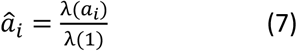

where λ(θ) is defined as in (2). Thus, λ(*a*_i_) expresses the sum of the observed matings having mutual-propensity *a*_i_, divided by the sum of the product of the population frequencies from each partner type. Similarly, λ(1) corresponds to the sum of the observed matings having unity mating parameter divided by the sum of the corresponding products of population frequencies (details in Appendix A).

#### 2.4.2 Intra-male competition

Relaxing condition (5-*ii*) implies that at least one male marginal propensity, say male of type *A*, is different from the rest of male types i.e. *m*_*Male_A*_ ≠ *m*_*Male_B*_ with *A* ≠ *B*. On the other side, the marginal propensity of females should be the same which means that there is no intra-female competition, all female types mate at an equal rate. The corresponding model can be obtained just by interchanging rows with columns in (6). Noting the parameters as *b*_j_ instead of *a*_i_, the maximum likelihood estimate is

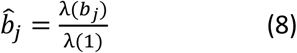

#### 2.4.3 Intra-female and male competition

By relaxing conditions (5-*i*) and (5-*ii*) the marginal propensities will be different within females and males. The corresponding model combines models (6) and (8) and has as much as (*k*_1_-1)*×*(*k*_2_-1) parameters in the most parameterized case, and a minimum of two (female and male) for the less parameterized one, in order to maintain the multiplicativity condition (5-*iii*). This type of model may produce a pattern of sexual selection in both sexes without assortative mating. By notational convenience, I fix the category *k*_1_ in females and *k*_2_ in males as having unitary parameters. Therefore

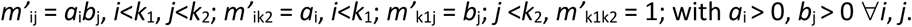

This model is multiplicative (see Appendix A) and the parameters MLE are

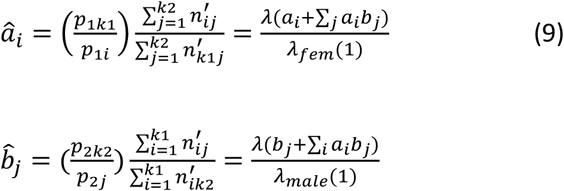

where the lambda function λ(*a*_i_ + *a*_i_*b*_1_ + … *a*_i_*b*_j_ + …) is applied to the mutual-propensities that depend on the parameter *a*_i_. Thus, λ(*a*_i_ +∑_j_ *a*_i_*b*_j_) is the quotient between the sum of the number of observed mating phenotypes that depend on the parameter *a*_i_ (i.e. ∑_j_ *n*’_ij_) and the sum of their expected random mating frequencies (which is simply *p*_1i_).

Correspondingly, λ_sex_(1) is the quotient between the sum of cases that contribute with 1 to the mutual-propensity by the given sex (i.e. ∑_j_ *n*’_k1j_ for females) and the sum of the expected frequencies (which is *p*_1k1_ for females). Formulae (9) is similar to (7) and (8). Note that the model in (9) becomes (7) by fixing every *b*_j_ as 1 while it becomes (8) by fixing every *a*_i_ as 1. The percentage of sexual selection information corresponding to each sex (*J*_S1_ and *J*_S2_ in Carvajal-Rodríguez 2018b), would depend on the population frequencies and on the mating parameter values.

### 2.5 Mate choice models

Mate choice models correspond to the class of non-multiplicative models, i.e. they can be obtained by relaxing the condition (5-iii) and may produce assortative mating patterns (positive or negative). If the female marginal propensities are equal and the same is true for the males (conditions 5-i and 5-ii hold) there would not be sexual selection neither in females nor males, and the model may produce only assortative mating patterns. However, this cannot be guaranteed in general because the occurrence of the sexual selection pattern is frequency dependent under non-multiplicative models (see below).

Consider a model where the unnormalized mutual-propensities are

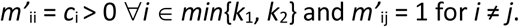

Thus, the homotype (*i*× *i*) mutual-propensities are parameterized while the heterotype are not. This model is non-multiplicative in general, because the contribution of the type *i* to the mutual-propensity is distinct in *m*_ii_ that in *m*_ij_ or in *m*_ji_ (although with an even number of types a multiplicative model can be obtained by setting *m*’_ii_ = 1 / *m*’_jj_).

By recalling the definition of marginal propensities in (4), the condition for equal female marginal *m*_Fem_i_ = *m*_Fem_j_ is

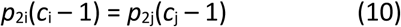

and in males

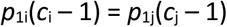

In general, depending on the conditions in (10), the mate choice models have double effect i.e. they produce assortative pattern jointly with sexual selection in at least one sex.

The maximum likelihood estimate for the model parameters is

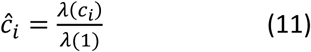

Note that the homotype mating parameter may imply higher mutual-propensity than the heterotype (*c*_i_ > 1, positive assortative mating) or viceversa, the homotype has lower mutual-propensity (*c*_i_ < 1, negative assortative). The number of different parameters ranges from 1 (*c*_1_ = *c*_2_ = … = *c*_i_) to *H*; where *H* = *min*{*k*_1_, *k*_2_} corresponds to the maximum possible number of different homotype matings.

It is also possible to define mate choice models with the heterotype mutual-propensities parameterized instead of the homotype ones (see Appendix A for details).

### 2.6 Models with mate competition and mate choice parameters

I have shown that mate choice models may generate both kinds of patterns, assortative mating and sexual selection, depending on the within sex population frequencies. While it is not possible to assure that the mate choice model produces no sexual selection, it is possible to combine the previous models to ensure that there are parameters directly linked to mate competition and parameters directly linked to mate choice. These combined models have the property that when the mate choice parameter is set to 1, there is only a known sexual selection effect caused by the competition parameter (female, male, or both). When the mate choice parameter is added, the assortative mating pattern appears and also, an extra effect of frequency-dependent sexual selection may be added to that of the original competition parameter.

#### 2.6.1 Models with male competition and mate choice: independent parameters

Consider the model *m*′_i1_ = *α*; *m*′_ii_ = *c* for *i* ≠ 1 and *m*′_ij_ = 1 otherwise; with *i*≤ *k*_1_, *j* ≤ *k*_2_. An example of this kind of model can be seen in Fig. 2.

**Fig 2.**
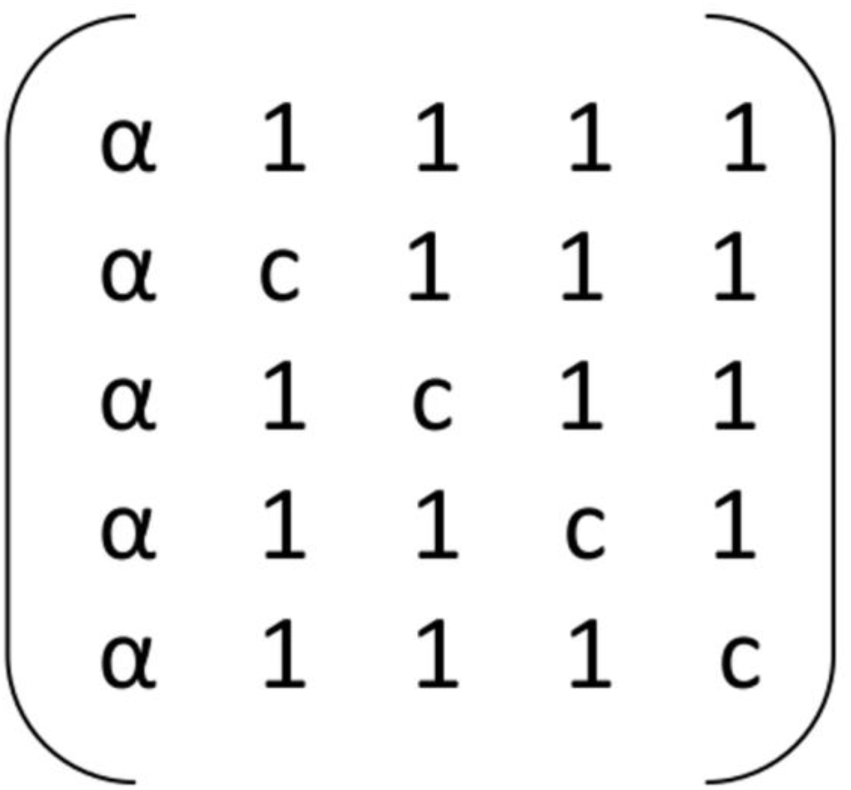
An example of male competition and mate choice independent parameters model with 5 × 5 mating phenotypes. α is the male competition parameter and *c* is the choice parameter. Rows are females, columns are males.

For the particular case of α ≠ 1, *c* = 1; the model has within male competition that corresponds to the marginal propensity α of the type-1 male compared with the other males, so, a male sexual selection pattern may be generated. On the contrary, the female marginal propensities are equal so there is no female competition. Considering mate choice and the assortative mating pattern, when *c* =1 the model is multiplicative so assortative mating should not occur. In fact, in this case the pair sexual isolation statistics (PSI) are equal (see Appendix A for details) and the assortative mating is 0, i.e., the overall index of sexual isolation *I*_*PSI*_ = 0 (*I*_*PSI*_ = (4ΣPSI_ii_ - Σ PSI_ij_)/(4ΣPSI_ii_ + Σ PSI_ij_)) (Carvajal-Rodríguez 2018b).

However, by taking *c* ≠ 1 a new component is added to the sexual selection pattern. The parameter *c* corresponds to mate choice and produces positive (*c* > 1) or negative (*c* < 1) assortative mating. The value of *I*_*PSI*_ is a function of the parameter *c* and the population frequencies. Female sexual selection may also emerge depending on the value of *c* and the population frequencies.

The MLEs of both parameters are

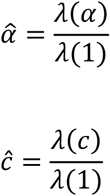

A variant of the above model can be generated by changing the *c* parameter from the main diagonal to the anti-diagonal. Similarly female sexual competition linked to the α-parameter is obtained by transposing the matrix of the model.

#### 2.6.2 Models with male competition and mate choice: compound parameters

Consider the model *m*′_11_ = *cα*; *m*′_i1_ = *α* and *m*′_ii_ = *c* for *i*>1; and *m*′_ij_ = 1 otherwise; with *i*≤*k*_1_, *j*≤*k*_2_. An example of this model can be seen in Fig. 3.

**Fig 3.**
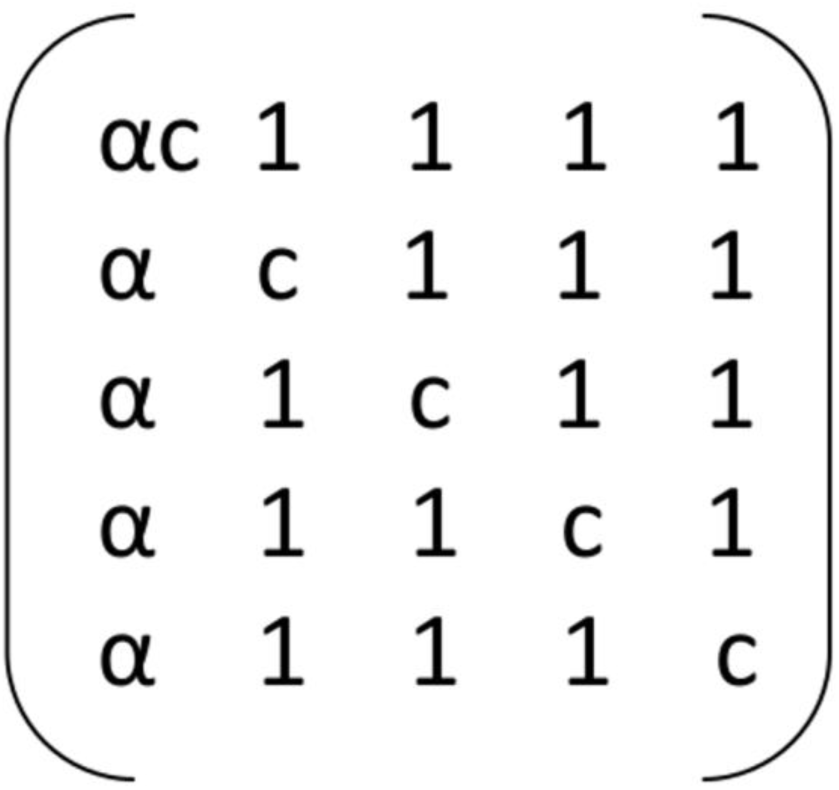
An example of male competition and mate choice compound parameters model with 5 × 5 mating phenotypes. α is the male competition parameter and *c* is the choice parameter. Rows are females, columns are males.

When *c* = 1 the model is the same as the previous one. When *c* ≠ 1, the mate choice parameter provokes an extra effect of sexual selection in males and females, plus assortative mating. The MLE of α and *c* are

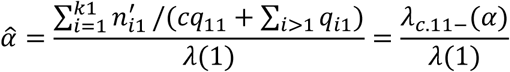

where λ_c.11-_(α) indicates that for matings with parameter α, the expected frequency indexed as 11 (i.e. q_11_) is weighted by *c*. Similarly,

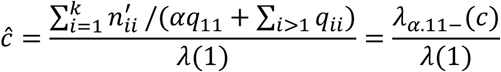

where *k* = min{*k*_1_, *k*_2_} and λ_α.11-_(*c*) indicates that for matings with mating parameter *c*, the expected frequency indexed as 11 is weighted by α.

The above estimates are dependent one on each other, so, for obtaining the estimates of this compound parameter model I have used a numerical bounded Nelder-Mead simplex algorithm, with restriction α > 0, *c* > 0 (Press 2002; Singer and Singer 2004; Gao and Han 2012).

#### 2.6.3 General model with male competition and mate choice parameters

The general model with male competition and mate choice parameters is *m*′_11_ = *c*_1_α; *m*′_i1_ = *α* and *m*′_ii_ = *c*_k_ for *i* > 1; and *m*′_ij_ = 1 otherwise; with *i*≤ *k*_1_, *j* ≤ *k*_2_. A particular case of this model can be seen in Fig. 4.

**Fig 4.**
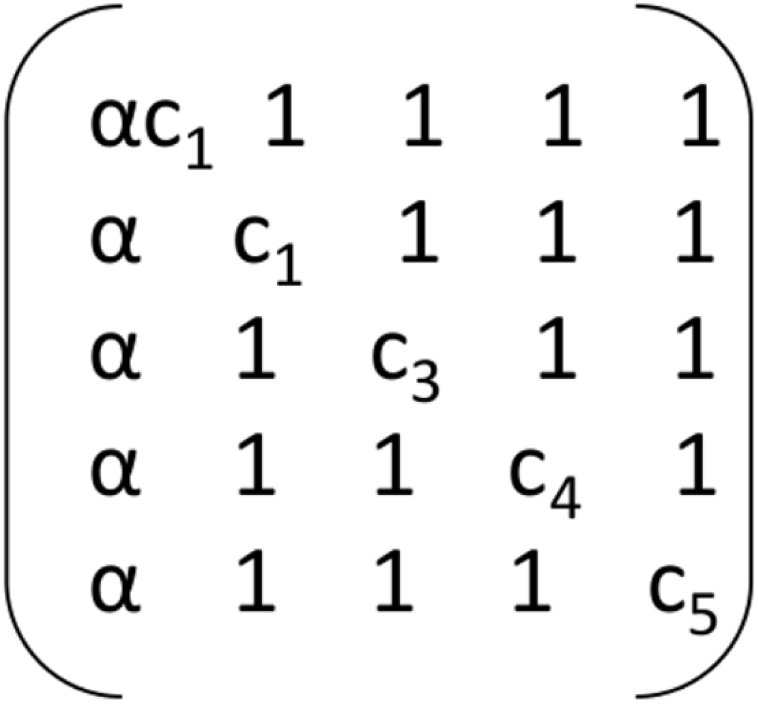
Male sexual selection and mate choice compound model with 5 × 5 mating phenotypes. α is the male sexual selection parameter and *c*_k_’s are the choice parameters with *c*_2_ = *c*_1_. Rows are females, columns are males.

Note that to distinguish the competition and mate choice parameters, it is necessary that at least one *c*_k_ parameter is equal to *c*_1_ (as in Figs. 3 and 4) or that *c*_1_ = 1 as in Fig. 2, otherwise the parameter for *m*_11_ does not distinguish competition and choice. Therefore, the model in Fig. 4 has *H* parameters with *H* = min{*k*_1_,*k*_2_} from which, *H - 1* are choice parameters (*c*) plus one male competition parameter α. The MLE are

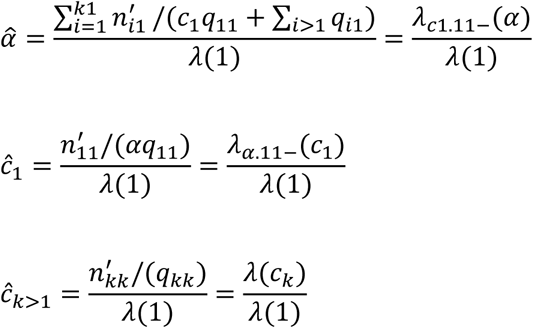

The model parameters *c*_k>1_ can be estimated directly from the sample; on the contrary, the α and *c*_1_ estimates are dependent on each other, so, for obtaining these estimates, I used a numerical bounded Nelder-Mead simplex algorithm with restriction α > 0, *c*_1_> 0 (Press 2002; Singer and Singer 2004; Gao and Han 2012).

Previous models were simplified versions of the general model. For example, the model in Fig. 2 is the general model with restrictions *c*_1_= 1; *c*_2_ = *c*_3_ =… = *c*_k_ = *c*. Also, the model in Fig. 3 corresponds to *c*_1_= *c*_2_ = *c*_3_ =… = *c*_k_ = *c*. Another particular case that could be defined is *c*_1_= *c*; *c*_2_ = *c*_3_ =… = *c*_k_ = 1. In the latter, the MLE of the parameters can again be expressed as a quotient of lambdas similar to the compound parameter case

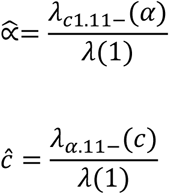

It is also possible to define another general model with the mate choice parameters in the anti-diagonal. Using the λ notation, the estimates follow the same formulae as defined for the general model with the choice parameters in the main diagonal. Concerning models with female competition and mate choice, they are obtained just by transposing the matrices of the mating parameters.

### 2.7 General double effect models

The mating parameters *m*_ij_ = θ_ij_ with the restriction that at least some are equal to one, permit to generate any particular model. In general, these models produce patterns of sexual selection and assortative mating with each parameter possibly linked to the occurrence of both (see Appendix A). The MLE is

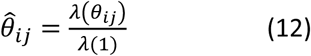

The most parameterized model of this kind is the saturated, with *K*-1 parameters. In such case, as already mentioned, the estimates in (12) are the corresponding pair total indices (PTI).

All the above derived MLE formulae have been verified by numerical approximation using the bounded Nelder-Mead simplex algorithm (Press 2002; Singer and Singer 2004; Gao and Han 2012).The set of described models jointly with their expected effects are summarized in Table 1.

**Table 1.**
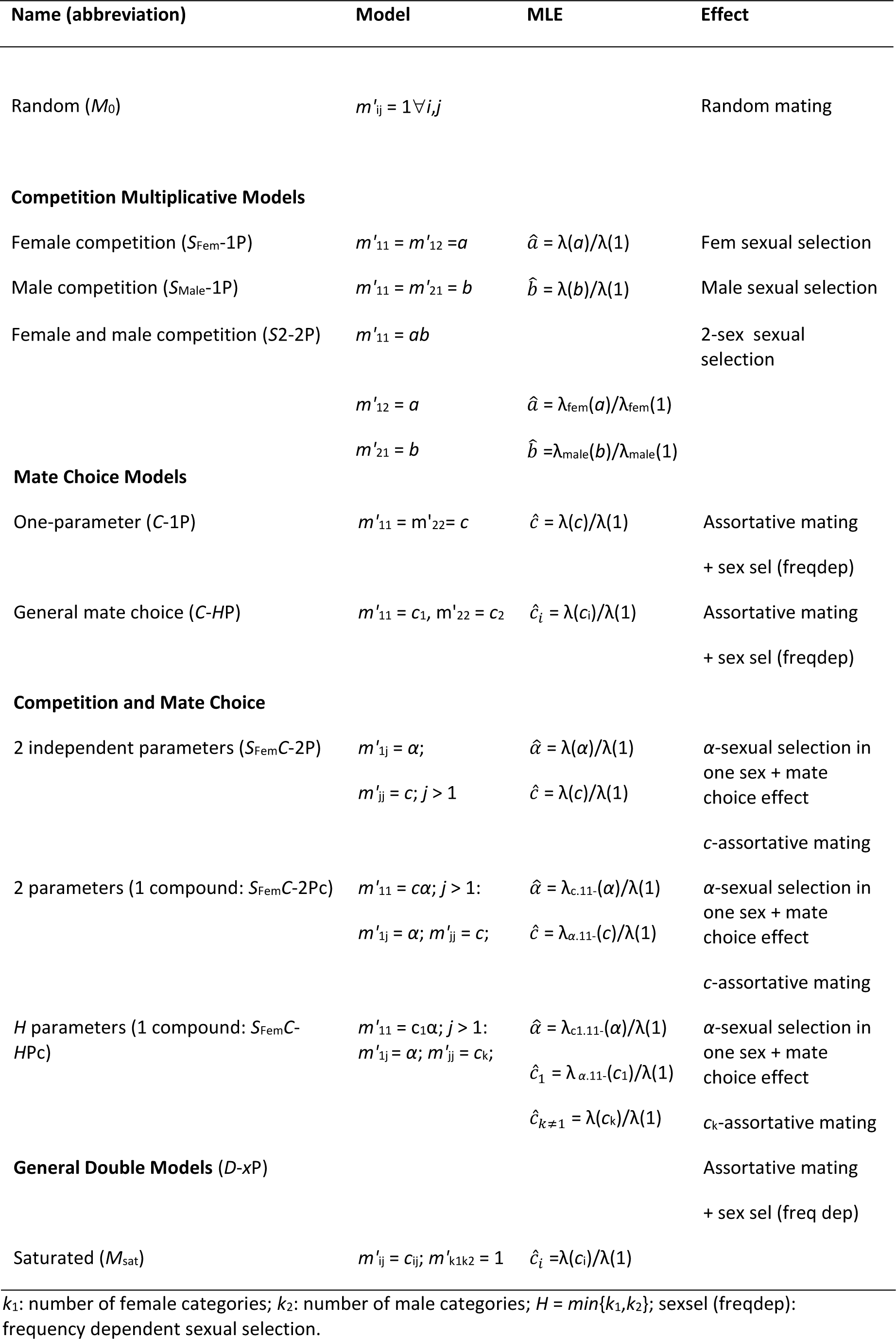
Mutual mating propensity models as defined by different parameters in a case with two different phenotypic classes in each sex (*k*_1_ = *k*_2_ = 2). The unnormalized *m*′_ij_ values not explicitly given are assumed to be 1.

## 3. Model selection and multi-model inference

Relying on the previous section, it would be possible to generate mate competition and mate choice models and, given a mating table, to apply some information criteria to select the best-fit candidates and estimating the mating parameter values based on the most supported models. Next, I briefly review the information criteria and model selection concepts and show how to apply them to perform model selection and multi-model inference among mate competition and mate choice models.

Information-based model selection and multi-model inference can be applied to describe uncertainty in a set of models to perform inference on the parameters of interest (Burnham et al. 2011; Grueber et al. 2011; Barker and Link 2015; Claeskens 2016). There are several information criteria at hand, although trusting on a single form of information criterion is unlikely to be universally successful (Liu and Yang 2011; Vrieze 2012; Brewer et al. 2016; Aho et al. 2017; Dziak et al. 2019). In the present work, two Kullback-Leibler divergence-based measures plus the so-called Bayesian information criterion are considered.

### 3.1. Information criteria

The Akaike information criterion (AIC) provides the link between the Kullback-Leibler divergence and the maximized log-likelihood of a given model (Akaike 1973). Here I use the sample-corrected version AICc, because it is asymptotically equivalent and may work better for small sample size

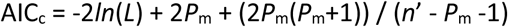

where *L* is the maximum likelihood of the model, *P*_m_ the total number of estimated mating parameters and *n*′ is the number of matings in the sample.

There is also a version for the symmetric Kullback-Leibler (Jeffrey′s) divergence, called the KICc criterion (Cavanaugh 2004; Keerativibool 2014). It seems adequate to consider the KICc criterion because the mating pattern obtained from the mutual-propensity models can be described by the informational flow from the mating frequencies, in the form of the Jeffrey′s divergence (Carvajal-Rodríguez 2018b) so,

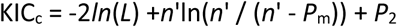

with *P*_2_ = *n*′[(*n*′ - *P*_m_)(2*P*_m_ + 3)-2] / [(*n*’ - *P*_m_ −2)(*n*′ - *P*_m_)]

Finally, the Bayesian information criterion (BIC Schwarz 1978) permits an approximation to the Bayes factor applied for model comparison (Wagenmakers 2007)

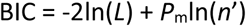

### 3.2 Overdispersion

In the context of model selection, data overdispersion, i.e. greater observed variance than expected, could generate the selection of overly complex models. The simplest approach to estimate overdispersion is by computing a single variance inflation factor (*v*). This inflation factor is the observed variation divided by the expected under the model with the highest likelihood (*M*_c_), other than the saturated, among the proposed ones (Richards 2008; Symonds and Moussalli 2011). It can be asymptotically approximated by the deviance i.e. twice the difference between the log-likelihood of the saturated (*M*_sat_) and the *M*_c_ model, divided by the difference in the number of parameters (*P*_Msat_ - *P*_Mc_) between both models

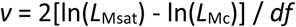

where *df* = *P*_Msat_ - *P*_Mc_.

If 1 ≤ *v* ≤ 4 this indicates overdispersion, while if higher than 4-6 this may indicate poor model structure and the construction of the set of models should be reconsidered (Burnham and Anderson 2002). For *v* values around 1 to 4, quasi-likelihood theory provides a way to analyse over dispersed data (Anderson et al. 1994; Richards 2008). The quasi-likelihood is the likelihood divided by an estimate of *v*. The quasi-likelihood version of the various information criteria, namely QAICc, QKIC_c_ (Kim et al. 2014) and QBIC, is obtained simply by replacing the likelihood with the quasi-likelihood in the corresponding formula. In such cases, the number of parameters is increased by one and the model variance is multiplied by *v* (see below). When the quasi-likelihood version is used, it must be done for all models and criteria.

### 3.3. Model weights

Let IC be any information criterion. For a particular criterion and for any set of *R* models there is a minimum criterion value e.g. AIC_cmin_, BIC_min_, etc. Thus, the models can be ranked regarding the difference with that minimum

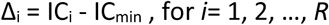

where IC_i_ refers to any specific information criterion for the model *i*.

Models can also be ranked by their weights from higher to lower. The weight *w*_i_ refers to the strength of evidence for that model (Burnham et al. 2011; Claeskens 2016)

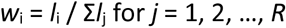

where *l*_i_ = exp(−0.5Δ_i_) is the relative likelihood of each model given the data.

### 3.4 Multi-model inference

Multi-model-based inference estimates the parameters of interest based on a group of the most credible models instead of on a best-fit single model (Burnham and Anderson 2002; Burnham et al. 2011; Symonds and Moussalli 2011). The multi-model inference is performed as a model averaged prediction for the parameters that are variables in the best model.

In our modelling framework and before performing the average of the estimated parameter values, the different models should be translated to the same scale of mutual-propensity. For example, a model like *m*′_11_ = 2, *m*′_12_ = *m*′_21_ = *m*′_22_ = 1, is not in the same scale that *m*′_11_ = 2, *m*′_12_ = *m*′_21_ = *m*′_22_ = 0.5. Without loss of generality, the latter can be transformed into an equivalent model *m*′_11_ = 4, *m*′_12_ = *m*′_21_ = *m*′_22_ = 1, which is now in the same scale as the first model.

The averaged parameter estimates were computed as a weighted mean where the weights are the strength of evidence for each model as obtained under a given information criterion. The parameters were averaged only over the models for which they appear as a variable. Because the weights need to sum up to 1, it was necessary renormalize them by dividing by the accumulated weight in the confidence subset.

Therefore, for each parameter *m* included in the confidence subset *R*_s_, the average was computed as

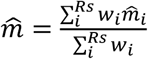

This way of performing the model averaged prediction is called natural averaging (Symonds and Moussalli 2011).

Finally, the reliability of each parameter estimate was measured as the unconditional standard error

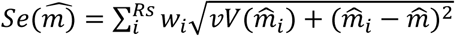

Where 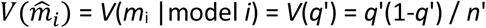 is the model standard error squared and *v* is the variance inflation factor.

The use of the sum of weights to estimate variable importance in regression models has been criticized because of multicollinearity among the predictor variables and the imprecision of the weight measures (Galipaud et al. 2014; Cade 2015; Galipaud et al. 2017). However, the mutual-propensity parameters do not belong to a regression model and their average is performed in the same scale and with comparable units. Therefore, under the mutual mating propensity setting, the multi-model inference would work well as it was confirmed by Monte Carlo simulation (next section).

## 4. Simulations

### 4.1 Polygamous species (sampling with replacement)

To test how well the above methodology is able to distinguish among the different classes of models and estimate the mating parameters, I used the sampling with replacement algorithm in the program MateSim (Carvajal-Rodríguez 2018a) to generate mating tables by Monte Carlo simulation (see Appendix B for detailed explanation).

The simulated cases correspond to one-sex competition and mate choice models. The resulting mating tables were consequence of the mating system and the sampling process, and consisted in two types of information (Fig. B1 in Appendix B). First, the population frequencies (premating individuals) which were generated randomly for each simulation run. Second, the sample of 500 mating pairs (*n*′ = 500) for a hypothetical trait with two classes at each sex. Because the simulated species had large population size (*n* = 10 000) the mating process was represented as a sampling with replacement, and the population frequencies were constant over the mating season. The minimum phenotype frequency (MPF) allowed was 0.1.

Five different model cases were simulated, namely random mating with mutual-propensities *m*′_11_=*m*′_22_ =*m*′_12_=*m*′_21_= 1 (*M*_0_ in Table 2), female competition (α = 2) and mate choice (*c* = 3) with independent parameters *m*′_11_= *m*′_12_ = 2, *m*′_22_ = 3, *m*′_21_ = 1 (SfC Table 2), and with compound parameters *m*′_11_= 6, *m*′_12_ = 2, *m*′_22_ = 3, *m*′_21_ = 1 (SfCc Table 2), and male competition (α = 2) and mate choice (*c* = 3) with independent parameters *m*′_11_= *m*′_21_ = 2, *m*′_22_ = 3, *m*′_12_ = 1 (SmC Table 2), and with compound parameters *m*′_11_= 6, *m*′_21_ = 2, *m*′_22_ = 3, *m*′_12_ = 1 (SmCc Table 2). Each case was simulated 1 000 times.

**Table 2.** Average (standard error) parameter estimates under sample size 500 for a polygamous species with large population size (*N* = 10 000).

| Model | | $m'_{11}$ | $m'_{12}$ | $m_{21}$ | $m_{22}$ |
| --- | --- | --- | --- | --- | --- |
| **M<sub>0</sub>** | **Expected** | **1** | **1** | **1** | **1** |
|  | **AICc** | 1.0 ± 0.0000 | 1.0 ± 0.0000 | 1.0 ± 0.0000 | 1.0 ± 0.0000 |
|  | **KICc** | 1.0 ± 0.0000 | 1.0 ± 0.0000 | 1.0 ± 0.0000 | 1.0 ± 0.0000 |
|  | **BIC** | 1.0 ± 0.0000 | 1.0 ± 0.0000 | 1.0 ± 0.0000 | 1.0 ± 0.0000 |
| **SfC** | **Expected** | **2** | **2** | **1** | **3** |
|  | **AICc** | 1.94 ± 0.0017 | 1.99 ± 0.0009 | 1.0 ± 0.0000 | 2.95 ± 0.0025 |
|  | **KICc** | 1.94 ± 0.0024 | 1.99±0.0020 | 1.0 ± 0.0000 | 2.95 ± 0.0029 |
|  | **BIC** | 1.90 ± 0.0060 | 1.94 ± 0.006 | 1.0 ± 0.0000 | 2.90 ± 0.0074 |
| **SfCc** | **Expected** | **6** | **2** | **1** | **3** |
|  | **AICc** | 5.93 ± 0.0044 | 2.0 ± 0.0001 | 1.0 ± 0.0000 | 2.97 ± 0.0027 |
|  | **KICc** | 5.92 ± 0.0051 | 2.0 ± 0.0001 | 1.0 ± 0.0000 | 2.97 ± 0.0028 |
|  | **BIC** | 5.87 ± 0.0086 | 2.0 ± 0.0017 | 1.0 ± 0.0000 | 2.96 ± 0.0044 |
| **SmC** | **Expected** | **2** | **1** | **2** | **3** |
|  | **AICc** | 1.94 ± 0.0020 | 1.0 ± 0.0000 | 1.99 ± 0.0013 | 2.94 ± 0.0030 |
|  | **KICc** | 1.93 ± 0.0032 | 1.0 ± 0.0000 | 1.98 ± 0.0029 | 2.93 ± 0.0037 |
|  | **BIC** | 1.90 ± 0.0062 | 1.00 ± 0.0000 | 1.93 ± 0.0065 | 2.88 ± 0.0080 |
| **SmCc** | **Expected** | **6** | **1** | **2** | **3** |
|  | **AICc** | 5.93 ± 0.0046 | 1.0 ± 0.0000 | 2.0 ± 0.0001 | 2.97 ± 0.0029 |
|  | **KICc** | 5.92 ± 0.0052 | 1.0 ± 0.0000 | 2.0 ± 0.0001 | 2.97 ± 0.0029 |
|  | **BIC** | 5.87 ± 0.0085 | 1.0 ± 0.0000 | 2.0 ± 0.0010 | 2.97 ± 0.0037 |

M<sub>0</sub>: Random mating model. SfC: female competition and mate choice with independent parameters. SfCc: female competition and mate choice with compound parameters. SmC: male competition and mate choice with independent parameters. SmCc: male competition and mate choice with compound parameters.

For each simulation run, and given the normalized mutual-propensities *m*_*ij*_, the number of occurrences for each mating class *i* × *j* was obtained as

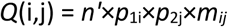

where *n*′ is the sample size, *p*_1i_ is the female population frequency for the phenotype *i, p*_2j_ is the male population frequency for the phenotype *j*.

Once the mating tables were obtained I proceeded with the multi-model inference analysis using InfoMating. Note that there were 1 000 different tables for each simulated case so, in the simulation study, it is better to consider the mean multi-model estimates instead of the full list of analysed models (which would imply 1 000 lists for each simulated case). Also, it is worth noting that with real data, the exactly true model is not necessarily included in the set of assayed models and so, it is important to evaluate the accuracy of the multi-model parameter estimates because, if the parameter estimates are correct, the model that would arise from that estimates and the set of most supported candidate models must be a good guess of the true one.

The sequence of analyses was as follows. For each mating table, InfoMating generates a set of 17 models, from the simplest random model M_0_ to the saturated M_sat_, including mate competition and choice models with one or two parameters (see all the types in Table 1). Then, the program computes the information criteria for each model and performs the multi-model inference as explained in the previous section. Thus, for each of the 5 simulated cases, 1 000 parameter estimates were obtained, and their average and standard error computed (Table 2).

It can be appreciated that the random mating was perfectly estimated by the three IC methods. The competition plus mate choice parameter estimates were fairly good under the three criteria. The estimates were slightly better under AICc and slightly less accurate under BIC.

The whole simulation process was repeated using a small sample size (*n*’ = 50 matings) and the results were qualitatively similar. However, the parameter estimates tended to be low-biased possibly because the power to detect deviations from random mating was low (see supplementary Table C1 in Appendix C).

### 4.2 Monogamous species (sampling without replacement)

For monogamous species, the mating process is without replacement (from the point of view of the available phenotypes) and can be represented via mass-encounters (Gimelfarb 1988; Carvajal-Rodríguez 2018a). The pattern obtained under the mass-encounter monogamous scenario (when the population size is large) was qualitatively similar to the polygamous species. However, there was less power to detect deviation from random mating and so the estimates were low-biased, especially in the case of the compound parameter. Regarding sample size, it seems that the estimation was not very much affected (see supplementary Tables C2 and C3 in Appendix C).

Not surprisingly, the case of monogamous species with small population size (*N* = 200) was the worst scenario for multi-model estimation under the assumption of constant population phenotype frequencies (see Table C4 in Appendix C). Under this case and when most of the adults were involved in the mating process (mating sample size = 100), the change in the population phenotype frequencies during the breeding season significantly affected the observed non-random mating patterns. Only when the deviation from random mating is as large as with the compound effect of choice and competition, the estimated mutual-propensities provided some information (SfCc in Table C4).

## 5. Example of application

*Littorina saxatilis* is a marine gastropod mollusc adapted to different shore habitats in Galician rocky shores. There are two different ecotypes, an exposed-to-wave (smooth un-banded, SU), and a non-exposed (rough banded, RB) ecotype. Several experimental studies have shown that these ecotypes have evolved local adaptation at small spatial scale. For example, stronger waves on the lower shore may provoke that the SU ecotype becomes sexually mature at smaller size than the upper-shore (RB) ecotype. In addition, in some areas of the mid-shore habitat, the two ecotypes occasionally mate, producing apparently fertile intermediate morphological forms that are called hybrids (HY) (Rolan-Alvarez et al. 2015a).

Sexual isolation (positive assortative mating) between RB and SU morphs was observed in wild mating pairs in the mid-shore zone, likewise within-morph size-assortative mating in all shore levels (Cruz et al. 2001). It is assumed that the size is the key trait causing the increase of sexual isolation in this model system, being the males the choosy sex in this species (Rolan-Alvarez 2007).

Here, I reanalysed a *L. saxatilis* data set (Cruz et al. 2001) to estimate the mutual-propensity parameters between the RB, SU and HY morphs in the mid-shore habitat. In the original study, the authors analysed a hybrid zone encompassing 30 km of coast in Galicia (NW Spain) with two sampling locations (Centinela and Senin) and seasons (autumn and summer). Mating pairs were collected jointly with the 15 nearest non-mating individuals. The classification of morphs was made by considering as pure morphs those snails that had their shell ridged and banded (RB morph) or smooth and unbanded (SU morph). The hybrids (HY) were those snails that had a complete set of bands but lacked ridges, or viceversa, or those that, having both ridges and bands, had at least two incomplete bands (see details in Cruz et al. 2001). In the present reanalysis, I considered the pooled data of the two sampling locations and seasons (Table 3).

**Table 3.** The population frequencies by sex and the sample of matings from Cruz et al. (2001) data.

|  | Total | RB | HY | SU |
| --- | --- | --- | --- | --- |
| **Female freqs** | 1254 | 0.22 | 0.11 | 0.67 |
| **Male freqs** | 1080 | 0.26 | 0.12 | 0.62 |
| **Matings** |  |  |  |  |
|  |  | **Males** |  |  |
|  |  | **RB** | **HY** | **SU** |
| **Females** |  |  |  |  |
|  | **RB** | 19 | 9 | 13 |
|  | **HY** | 6 | 4 | 10 |
|  | **SU** | 6 | 7 | 80 |

First, I computed the information partition (Carvajal-Rodríguez 2018b) that indicated significant assortative mating from the Chi-square test (*J*_PSI_ *p*-value < 0.0000001) while no significant sexual selection was detected. However, the randomization test was not significant in any case, possibly due to the low sample size within the mating classes. Second, I proceeded with the model estimation and initially assayed only the subset of models with male and/or female mate competition plus the saturated (*M*_sat_) and random mating (*M*_0_) models. The estimate of overdispersion was high (7.20) indicating poor structure of the set of models regarding the data. The three information criteria gave similar output with the *M*_0_ as the best fit model. The multi-model estimates of the mutual-propensities were just one in every case as expected from random mating. Because in the simulation study, the AICc criterion gave the best estimates I will rely on this criterion from now on.

The next step was to study only models with choice parameter plus the saturated (*M*_sat_) and random mating (*M*_0_) models. The overdispersion was 4.65 that still indicates somewhat poor model structure. The best fit model was a choice model with one parameter. The multi-model inference gave a clear pattern of positive assortative mating, that was higher for the RB × RB mating (*m*’_RBRB_ = 3), intermediate for HY × HY (*m*’_HYHY_ = 2.3) and slightly lower for SU × SU (*m*’_SUSU_ = 2).

Then, I considered jointly the previous competition and choice models and added new ones having separated competition and choice parameters. The overdispersion was 3.4 that is an acceptable value for multinomial models and can be corrected by using quasi-likelihoods (see the overdispersion section above). Now, the best fit was a compound parameter model with female competition and choice. The estimates from this model were a RB female competition of α = 1.7 and choice *c* = 2.4. The multi-model estimates gave positive asssortative mating, *m*’_RBRB_ = 3, *m*’_HYHY_ = 3, *m*’_SUSU_ = 2 and sexual selection favouring RB females.

Finally, I considered all the previous models plus models having parameters with double effect (i.e. one parameter may generate both sexual selection and assortative mating patterns). This implies a total of 35 models including *M*_0_ and *M*_sat_. The overdispersion was 2.5. The best model was the same for the three criteria and it was a double effect model with 2 parameters, *c*_1_ = 0.2 and *c*_2_ = 2, distributed as indicated in Fig. 5. Approximately, the same model was obtained using the multi-model estimates.

**Fig 5.**
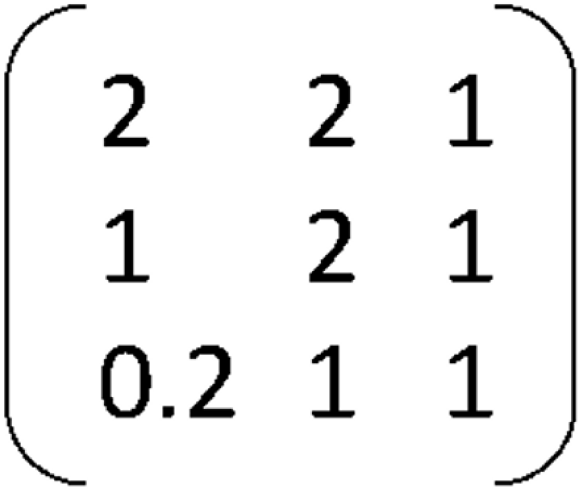
Model D-2P-Rep3: Double two parameter model with three repetitions of the *c*_2_ parameter (*c*_1_ = 0.2, *c*_2_ = 2) producing female and male sexual selection plus positive assortative mating.

It is also possible to focus only on the models with separated parameters for competition and choice. The best fit model from this subgroup involves female competition. Recall that in *Littorina saxatilis* the choosy sex are the males, so I considered that the competitive advantage from the side of the females is explained by the males preferring a given kind of females. The best fit model is *S*_Fem_*C*-2Pc (see Table 1) with RB female competitive advantage of 1.7 more times matings than the other females and a choice parameter of 2.4. The qualitative pattern obtained from these models is similar to that in Fig.5; the RB females (first row) are preferred and there is a choice for within ecotype mating. The combination of competition and choice provokes that the mating RB×RB is the preferred by RB males (first column in Fig. 5), the matings RB×HY and HY×HY are preferred by HY males (second column in Fig. 5), and finally, it seems that the SU males (third column in Fig. 5) do not discriminate between female ecotypes.

## 6. Discussion

### 6.1 Simulations

I have simulated mating tables corresponding to random mating, mate competition and mate choice models. The random mating pattern was perfectly assessed. For the other models, the competition and choice parameters were estimated quite accurately when the mating system resembles a sampling with replacement. Not surprisingly, BIC was slightly more conservative, while AICc presented slightly more accurate estimates in most cases. The KICc criterion performed similar to the best AICc and BIC cases. In general, the estimation was accurate and even in the cases with extreme phenotypic frequencies, the mean estimates were closer to the real value than to random mating.

The proposed approach does only require mating tables. However, to correctly identify the processes that produce the patterns of sexual selection and assortative mating, it is assumed that the encounters occur at random, i.e. the encounter between two phenotypes depends on the population phenotypic distribution, and that the mating pattern is the product of the phenotypic distribution of the population and the individual preferences (Carvajal-Rodríguez 2018a). As a consequence, the availability of phenotypes should not be affected by the matings that have already occurred, as expected for polygamous species, or even for monogamous species, when the number of available individuals is higher than the mating pairs.

However, the above assumption is likely to be violated in the case of monogamous species with low population size, or even in large population sizes with local competition for mates (if the number of individuals in the patches is low) and/or space-temporal constraints. In such cases, the mating process resembles a sampling without replacement and the population phenotype frequencies may be altered during the reproductive season so that the sexual selection and assortative mating patterns would be more difficult to detect (Carvajal-Rodríguez 2019). In fact, the simulations (see Appendix C) showed that the performance of the multi-model inference is affected by the sampling and the mating system (polygamous or monogamous) but it is still quite robust for detecting non-random mating in the parameter values except in the worst scenario of monogamous species with small population sizes.

### 6.2 General

The advantages of model selection and multi-model inference in evolutionary ecology has been widely discussed, jointly with the pros and cons of applying any information criteria (Link and Barker 2006; Burnham et al. 2011; Aho et al. 2014; Barker and Link 2015; Aho et al. 2017; Dziak et al. 2019) or the reliability of the obtained estimates (Galipaud et al. 2014; Cade 2015; Giam and Olden 2016; Galipaud et al. 2017).

Multi-model inference has been however, rarely utilized to study the mating patterns that may emerge from mate choice and mate competition. Here, by developing general models that incorporate competition and mate choice, and providing their maximum likelihood estimates, I am proposing a standardized methodology for model selection and multi-model inference of the mating parameters producing the sexual selection and assortative mating patterns.

The set of *a priori* models permits to perform an *a posteriori* quantification of the data-based evidence and provide confidence sets on plausible non-trivial models while letting multi-model inference of the parameter values. The approach was implemented by allowing three different information criteria. Under the scenarios assayed, they performed similarly for simulated and real data.

Regarding the methodology, it is worth noting that although the mating tables require at least two phenotypes by sex (2×2 dimensions or higher) for fitting mate competition and mate choice models, the proposed approach can still be applied if some sex, say females, have only one phenotypic class. In this case, we just need to duplicate the row (see Fig. D1 in the Appendix D). Obviously, there cannot be any assortative pattern and sexual selection can only be measured in the sex with more than one phenotypic class.

The statistical tools developed in this work have been also applied to empirical data. Previous studies in the Galician *L. saxatilis* hybrid zone showed that mate choice favours within-morph pairs (reviewed in Rolan-Alvarez 2007). The estimates obtained by multi-model inference support the positive assortative mating for the ecotype. In addition, another result emerged from the analysis: The RB females are preferred in general i.e. RB male with SU female has less mutual-propensity than SU male with RB female (*m*_SURB_ < *m*_RBSU_). This pattern may be favoured by the physical difficulty for the mating involving bigger RB males with the smaller SU females, and could be related with the somehow more frequent occurrence of mating pairs having females bigger than males (a typical trend in gastropods, E. Rolán-Alvarez personal communication). Besides the mating pattern depicted by the multi-model approach, the estimates of the mutual-propensities were also obtained. Testing the reliability of these estimates is, however, out of the scope of the present manuscript, and it was left for future work.

To conclude, I present a methodology to distinguish among several models of mate competition and choice behind the observed pattern of mating and the phenotypic frequencies in the population. From an empirical point of view it is much easier to study patterns than processes and this is why the causal mechanisms of natural and sexual selection are not so well known as the patterns they provoke. I propose a new tool that will help to distinguish among different alternative processes behind the observed mating pattern.

## 7. Software, code and data accessibility

The developed methodology has been fully implemented in a program called InfoMating available at http://acraaj.webs6.uvigo.es/InfoMating/Infomating.htm or upon request to the author. The simulations data set is available at: https://doi.org/10.5281/zenodo.2749692

## Acknowledgements

I would like to thank Carlos Canchaya, Emilio Rolán-Alvarez, Sara Magalhaes, Alexandre Courtiol and two anonymous reviewers for many valuable comments that significantly improved this paper. E. R-A also provided the data for the example. This work was supported by Xunta de Galicia (Grupo de Referencia Competitiva, ED431C2016-037), Ministerio de Economía y Competitividad (CGL2016-75482-P) and by Fondos FEDER (“Unha maneira defacer Europa”). This preprint has been peer-reviewed and recommended by Peer Community In Evolutionary Biology (https://doi.org/10.24072/pci.evolbiol.100075)

## APPENDIX

### Appendix A) Mutual Mating Propensity Models

#### Saturated non-random mating model: λ notation

Consider the total number of possible mating phenotypes *K* = *k*_1_ × *k*_2_ and the saturated multinomial model for the *K*-1 free mating parameters *m*′_ij_.

The log-likelihood function is

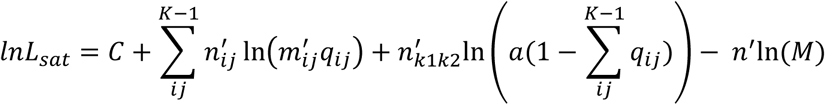

where *n*′ is the number of matings in the sample and *n*′_ij_ is the number of matings between *i*-type females and *j*-type males. I have fixed the parameter *m*′_k1k2_ to *a*.

Compute the first derivative of the likelihood with respect to *a*

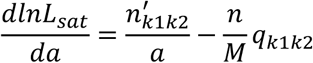

then by taking *a* = 1 and equating to 0 we get

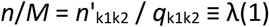

that corresponds to the number of observed matings having unity mating parameter divided by the corresponding product of population frequencies. Under the saturated model there is only one (for convenience *m*′_k1k2_) mating parameter having unitary value and so the number of observed matings is *n*′_k1k2_ and the product of the corresponding population frequencies is *p*_1k1_ × *p*_2k2_ = *q*_k1k2_.

Now, let find the *m*′_ij_ parameter value that maximizes the likelihood

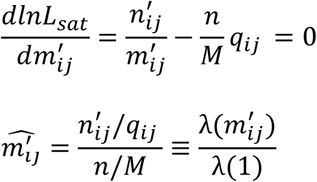

The λ notation can be generalized for any set *A* of mating phenotypes having the same value of propensity θ as follows

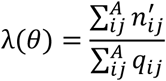

where *n*’_ij_ represents the number of mating pairs having absolute (non-normalized) mating parameter θ and *q*_ij_ is the product of the population frequencies *p*_1i_ and *p*_2j_ i.e. the expected frequency of the θ-mating phenotypes under random mating.

#### Intrafemale competition models

The model is

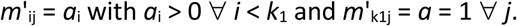

There are *k*_1_-1 independent parameters. Note that the parameters *m*′_k1j_ have been fixed to *a* = 1. The log-likelihood function is

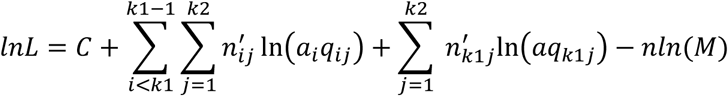

Now, assume that the parameter *a* is not fixed and compute the first derivative of the likelihood with respect to *a*

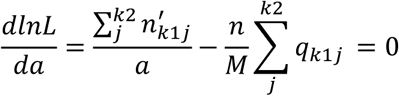

then by taking *a* = 1 and equating to 0 we get

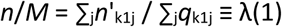

Now find the *a*_i_ parameter value that maximizes the likelihood

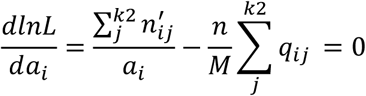

Solving for *a*_i_

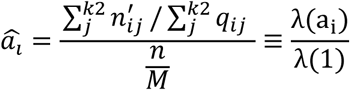

The formula expressed as the quotient of lambdas is valid for any number *h* of different parameters, 1 ≤ *h* < *k*_1_. In the particular case of having only one parameter the sum of observed matings having propensity *a*_1_, implies ΣΣ*n*′_ij_ where the first summation is for all the female types except females of type *k*_1_, and the second is over all male types. The sum of the product of frequencies is 1 -*p*_1k1_.

As before, λ(1) also corresponds to the sum of the observed matings having expected propensity 1 divided by the sum of the corresponding products of population frequencies. The model for male sexual selection is solved in a similar way.

#### Intrasexual competition in both sexes

The model is

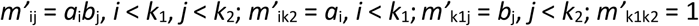

with *a*_i_ > 0, *b*_j_ > 0 ∀ *i, j*.

It is easy to see that is multiplicative. Let 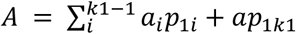 and 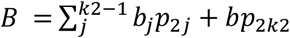

The mean mutual mating propensity is

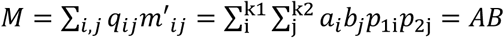

with *a*_k1_ = *a* and *b*_k2_ = *b*.

The marginal propensity for *i*-type females is

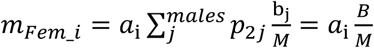

Similarly, the marginal for *j*-type males

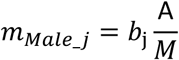

with *a*_k1_ = *a* and *b*_k2_ = *b*.

The condition 5-iii) for a multiplicative model implies that *m*_ij_ = *m*_Fem_i_ × *m*_Male_j_. In addition, *m*_ij_ = *a*_i_*b*_j_/*M* that jointly with the multiplicative condition requires *a*_i_*b*_j_/*M* = *m*_Fem_i_ × *m*_Male_j_ = *a*_i_*Bb*_j_*A* /*M*^2^ solving for *M* we get *M* = *AB* which we have already seen it is true.

The log-likelihood function

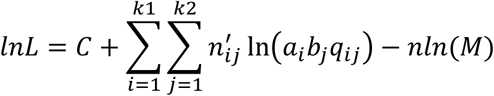

with *a*_k1_ = *a* = 1 and *b*_k2_ = *b* = 1.

Consider the derivatives

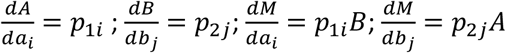

Now by taking the derivative of the log-likelihood with respect to *a*_i_ or *b*_j_ and equating to 0 we get the estimates

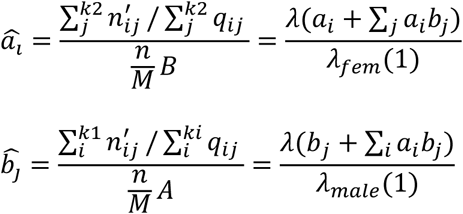

Where

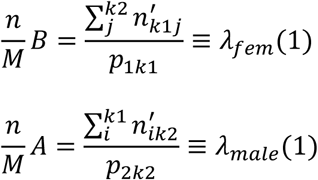

#### Mate choice models with parameterized heterotypes

Consider models in which the homotype mating has absolute propensity of 1 while the different heterotypes have absolute value of *c*_ij_. The maximum likelihood estimate is

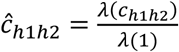

The number of parameters in this type of model is *K* – *min*{*k*_1_, *k*_2_} – Σ_S_(C_s_-1) where the sum is over the set of different heterotype matings and C_s_ is the cardinality of each set.

#### Double effect models

The following models generate a double pattern of sexual selection and assortative mating even when the population frequencies are uniform.

#### Double effect models producing sexual selection in one sex under uniform frequencies

A simple approach consists in building a new model by setting *m*′_ii_ = 1 and *m*′_jj_ = 1 + *c*. Then, if we desire assortative mating jointly with sexual selection only in females we additionally set *m*′_ij_ = 1 – *c*; on the contrary, if we desire selection only in males we set *m*′_ji_ = 1 – *c* with −1 < *c* < 1. If the frequencies are not uniform the model generates assortative mating jointly with sexual selection in both sexes.

In the case of the model with *m*′_ij_ = 1 -*c* (female sexual selection if frequencies are uniform) the maximum likelihood estimate of *c* is one of the roots of the quadratic

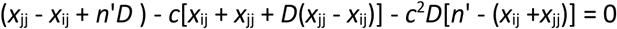

where *D* = *q*_ij_ - *q*_jj_ and *n*′ = ∑*x*_ij_ is the number of matings (sample size).

If the frequencies are uniform and *k*_1_ = *k*_2_, i.e. *p*_1i_ = *p*_1j_ = *p*_2i_ = *p*_2j_ ∀ *i, j* then

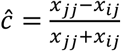

The case for male sexual selection is obtained simply by interchanging *x*_ij_ by *x*_ji_ and *q*_ij_ by *q*_ji_ in the formulas.

The above model has only one parameter *c*; we can introduce a more complex two parameter model, *M*_(a,c)_ by setting *m*′_ii_ = *a, m*′_jj_ = 1 +*c* and *m*′_ij_ = 1 - *c*, for female sexual selection (or *m*′_ji_ = 1 - *c* for male sexual selection). For obtaining the MLE of this two parameter double model, with restrictions *a* > 0, *c* < |1|, I have used a numerical bounded Nelder-Mead simplex algorithm (Press 2002; Singer and Singer 2004; Gao and Han 2012).

#### Double effect models with sexual selection in both sexes under uniform frequencies

To get assortative mating jointly with sexual selection in both sexes under uniform frequencies, we just need to combine the above uniform one parameter models of each sex, so that *m*′_ii_ = 1, *m*′_jj_ = 1 +*c* and *m*′_ij_ = *m*′_ji_ = 1 - *c*.

The maximum likelihood estimate of *c* involves the solution of the quadratic

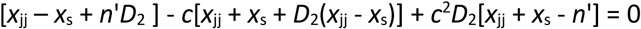

where *x*_s_ = *x*_ij_ + *x*_ji_ and *D*_2_ = *q*_ij_ + *q*_ji_ –*q*_jj_.

#### General double effect models

We can also define a set of general models where any propensity *m*′_ij_ has parameter θ_ij_ with at least one propensity having value of 1. The MLE of the parameters of this kind of model is

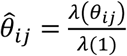

where λ(θ_ij_) is defined as in (A2).

The simplest model defined in this way is

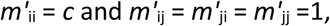

which produces assortative mating and sexual selection in both sexes.

Consider as an example of this model, the case with *k*_1_ = *k*_2_ = 2 so that 0 < *p*_11_ < 1; 0 < *p*_21_ < 1; *m*′_11_ = *c* and *m*′_12_ = *m*′_21_ = *m*′_22_ =1. The mean mating propensity is *M* = *q*_11_(*c* - 1) +1. The absolute marginal propensity for the first female type *m’*_*Fem_1*_ = *cp*_21_ + 1 – *p*_21_ = *p*_21_(*c* - 1) +1, and for the second female type *m’*_*Fem_2*_ = 1. Similarly the male marginals are *m’*_*Male_1*_ = *p*_11_(*c* - 1) +1 and *m’*_*Male_2*_ = 1.

Recall that the condition for the sexual selection pattern within a given sex is that the marginal mating propensities are different which here is true for both sexes provided that *c* ≠ 1. Regarding the assortative mating pattern it can be proved that the joint isolation index (I_PSI_) is 0 only if *c* = 1. However, it is sufficient to prove that the model is not multiplicative (Carvajal-Rodríguez 2018b). Consider that the model is multiplicative, this implies, *m*′_12_ /*M* = (*m’*_*Fem_1*_ /*M*) × (*m’*_*Male_2*_ /*M*) that given the model values becomes

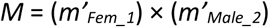

which can be true only when *p*_11_ = 1 and so it is false by definition.

The estimate of *c* under this model is λ(*c*) / λ(1).

The most parameterized model that can be defined in this way has *K*-1 free parameters and coincides with the saturated model so that the estimates are the corresponding pair total indices (PTI_ij_).

Moreover, note that if no mutual propensity is fixed to 1 then λ(1) = (*n* - *A*) / (1-*P*) = *n* where *A* = number of observations having value 1 = 0 and *P* = product of population frequencies of the involved types having mutual propensity 1 = 0. Therefore the estimate of θ_ij_ can also expressed as λ(θ_ij_) /*n* which is the observed frequency of mating pairs (*i, j*) divided by the expected frequency by random mating which is the definition of the pair total index PTI_ij_ (*K*-1 are free and one PTI is dependent on the others).

All the above derived MLE formulae have been checked by a numerical bounded Nelder-Mead simplex algorithm (Press 2002; Singer and Singer 2004; Gao and Han 2012).

### Appendix B) Monte Carlo simulation of mating tables

The mating tables for the simulation experiments were generated by the program MateSim (Carvajal-Rodríguez 2018a) available at http://acraaj.webs.uvigo.es/MateSim/matesim.htm. The number of replicates for each case was 1 000. For each run the program first generated the number of premating males and females from a given population size. For example, if the population size consisted in *n*_1_ (= 5 000) females and *n*_2_ (= 5 000) males, the program got *n*_1A_ = *n*_1_ × *U* females of the A type and *n*_1B_ = *n*_1_ - *n*_1A_ females of the B type. Where *U* is a value sampled from the standard uniform distribution. The premating males were obtained similarly. Then, the female population frequencies were *p*_1i_ = *n*_1i_ / *n*_1_, and *p*_2i_ = *n*_2i_ / *n*_2_ for the male ones. Finally, a sample of *n*′ (= 500) matings was obtained, where the number of counts for each mating phenotype *i* × *j* was

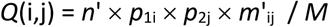

where *m*′_ij_ are the mutual-propensity parameters as defined for each kind of model, and *M* 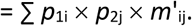.

The format of the obtained tables was the same as the JMating (Carvajal-Rodriguez and Rolan-Alvarez 2006) input files (Fig. B1).

**Fig. B1.**
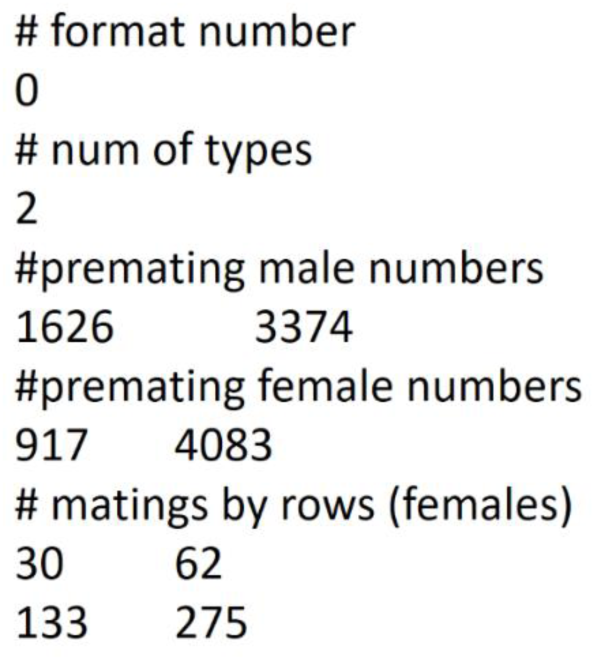
Example of a table generated by the simulations. The format is the same as for the JMating software.

### Appendix C) Polygamous species with low sample size and monogamous species

**Table C1.**
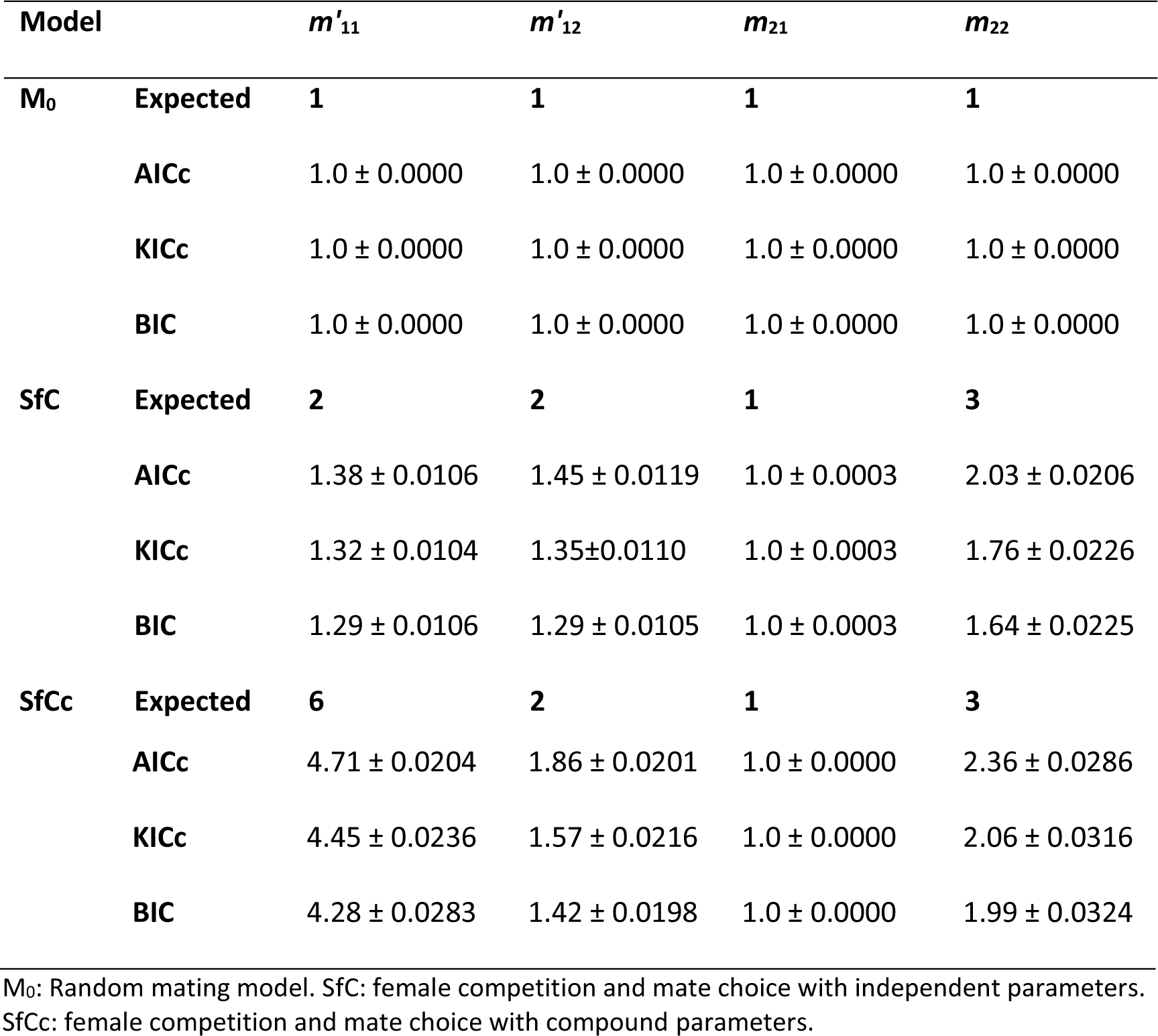
Average (standard error) parameter estimates under sample size 50 for a polygamous species with large population size (*N* = 10 000).

| Model | | $m'_{11}$ | $m'_{12}$ | $m_{21}$ | $m_{22}$ |
| --- | --- | --- | --- | --- | --- |
| **M<sub>0</sub>** | **Expected** | **1** | **1** | **1** | **1** |
|  | **AICc** | 1.0 ± 0.0000 | 1.0 ± 0.0000 | 1.0 ± 0.0000 | 1.0 ± 0.0000 |
|  | **KICc** | 1.0 ± 0.0000 | 1.0 ± 0.0000 | 1.0 ± 0.0000 | 1.0 ± 0.0000 |
|  | **BIC** | 1.0 ± 0.0000 | 1.0 ± 0.0000 | 1.0 ± 0.0000 | 1.0 ± 0.0000 |
| **SfC** | **Expected** | **2** | **2** | **1** | **3** |
|  | **AICc** | 1.38 ± 0.0106 | 1.45 ± 0.0119 | 1.0 ± 0.0003 | 2.03 ± 0.0206 |
|  | **KICc** | 1.32 ± 0.0104 | 1.35±0.0110 | 1.0 ± 0.0003 | 1.76 ± 0.0226 |
|  | **BIC** | 1.29 ± 0.0106 | 1.29 ± 0.0105 | 1.0 ± 0.0003 | 1.64 ± 0.0225 |
| **SfCc** | **Expected** | **6** | **2** | **1** | **3** |

|  |  |  |  |  |
| --- | --- | --- | --- | --- |
| **AICc** | $4.71 \pm 0.0204$ | $1.86 \pm 0.0201$ | $1.0 \pm 0.0000$ | $2.36 \pm 0.0286$ |
| **KICc** | $4.45 \pm 0.0236$ | $1.57 \pm 0.0216$ | $1.0 \pm 0.0000$ | $2.06 \pm 0.0316$ |
| **BIC** | $4.28 \pm 0.0283$ | $1.42 \pm 0.0198$ | $1.0 \pm 0.0000$ | $1.99 \pm 0.0324$ |

M<sub>0</sub>: Random mating model. SfC: female competition and mate choice with independent parameters. SfCc: female competition and mate choice with compound parameters.

**Table C2.** Average (standard error) parameter estimates under sample size 500 for a monogamous species (mass-encounter mating process) with large population size (*N* = 10000).

| Model | | $m'_{11}$ | $m'_{12}$ | $m_{21}$ | $m_{22}$ |
| --- | --- | --- | --- | --- | --- |
| **M<sub>0</sub>** | **Expected** | **1** | **1** | **1** | **1** |
|  | **AICc** | 1.07 ± 0.0053 | 1.07 ± 0.0053 | 1.07 ± 0.0059 | 1.07 ± 0.0056 |
|  | **KICc** | 1.05 ± 0.0044 | 1.04 ± 0.0043 | 1.05 ± 0.0052 | 1.04 ± 0.0050 |
|  | **BIC** | 1.01 ± 0.0028 | 1.01 ± 0.0020 | 1.01 ± 0.0033 | 1.01 ± 0.0026 |
| **SfC** | **Expected** | **2** | **2** | **1** | **3** |
|  | **AICc** | 1.95 ± 0.0158 | 1.95 ± 0.0160 | 1.0 ± 0.0001 | 2.04 ± 0.0176 |
|  | **KICc** | 1.93 ± 0.0162 | 1.93 ± 0.0166 | 1.0 ± 0.0001 | 2.03 ± 0.0180 |
|  | **BIC** | 1.82 ± 0.0183 | 1.82 ± 0.0189 | 1.0 ± 0.0000 | 1.93 ± 0.0195 |
| **SfCc** | **Expected** | **6** | **2** | **1** | **3** |
|  | **AICc** | 3.02 ± 0.0341 | 2.07 ± 0.0210 | 1.0 ± 0.0001 | 2.84 ± 0.0298 |
|  | **KICc** | 3.0 ± 0.0343 | 2.04 ± 0.0216 | 1.0 ± 0.0000 | 2.82 ± 0.0302 |
|  | **BIC** | 2.86 ± 0.0353 | 1.92 ± 0.0244 | 1.0 ± 0.0000 | 2.73 ± 0.0313 |

M<sub>0</sub>: Random mating model. SfC: female competition and mate choice with independent parameters. SfCc: female competition and mate choice with compound parameters.

**Table C3.** Average (standard error) parameter estimates under sample size 50 for a monogamous species (mass-encounter mating process) with large population size (*N* = 10000).

| Model | | $m'_{11}$ | $m'_{12}$ | $m_{21}$ | $m_{22}$ |
| --- | --- | --- | --- | --- | --- |
| **M<sub>0</sub>** | **Expected** | **1** | **1** | **1** | **1** |
|  | **AICc** | 1.0 ± 0.0362 | 1.04 ± 0.0414 | 1.0 ± 0.0365 | 1.02 ± 0.0406 |
|  | **KICc** | 1.02 ± 0.0339 | 1.03 ± 0.0374 | 1.0 ± 0.0324 | 1.02 ± 0.0377 |
|  | **BIC** | 1.03 ± 0.0312 | 1.03 ± 0.0340 | 1.0 ± 0.0283 | 1.03 ± 0.0361 |
| **SfC** | **Expected** | **2** | **2** | **1** | **3** |
|  | **AICc** | 2.0 ± 0.0568 | 2.05 ± 0.0638 | 1.08 ± 0.0155 | 2.23 ± 0.0635 |
|  | **KICc** | 1.76 ± 0.0539 | 1.78 ± 0.0605 | 1.06 ± 0.0145 | 1.96 ± 0.0597 |
|  | **BIC** | 1.62 ± 0.0515 | 1.66 ± 0.0578 | 1.05 ± 0.0143 | 1.79 ± 0.0555 |
| **SfCc** | **Expected** | **6** | **2** | **1** | **3** |
|  | **AICc** | 3.36 ± 0.1004 | 2.19 ± 0.0731 | 1.03 ± 0.0073 | 3.11 ± 0.0997 |
|  | **KICc** | 3.06 ± 0.0951 | 1.94 ± 0.0715 | 1.03 ± 0.0066 | 2.85 ± 0.0964 |
|  | **BIC** | 2.89 ± 0.0919 | 1.79 ± 0.0687 | 1.02 ± 0.0048 | 2.68 ± 0.0929 |

M<sub>0</sub>: Random mating model. SfC: female competition and mate choice with independent parameters. SfCc: female competition and mate choice with compound parameters.

**Table C4.** Average (standard error) parameter estimates under sample size 100 for a monogamous species (mass-encounter mating process) with small population size (*N* = 200).

| Model | | $m'_{11}$ | $m'_{12}$ | $m_{21}$ | $m_{22}$ |
| --- | --- | --- | --- | --- | --- |
| **M<sub>0</sub>** | **Expected** | **1** | **1** | **1** | **1** |
|  | **AICc** | 1.07 ± 0.0083 | 1.10 ± 0.0141 | 1.09 ± 0.0141 | 1.07 ± 0.0090 |
|  | **KICc** | 1.04 ± 0.0064 | 1.05 ± 0.0094 | 1.05 ± 0.010 | 1.04 ± 0.0070 |
|  | **BIC** | 1.02 ± 0.0047 | 1.02 ± 0.0056 | 1.02 ± 0.0065 | 1.02 ± 0.0052 |
| **SfC** | **Expected** | **2** | **2** | **1** | **3** |
|  | **AICc** | 1.15 ± 0.0173 | 1.11 ± 0.0157 | 1.07 ± 0.0118 | 1.11 ± 0.0145 |
|  | **KICc** | 1.10 ± 0.0141 | 1.06 ± 0.0125 | 1.04 ± 0.0091 | 1.06 ± 0.0108 |
|  | **BIC** | 1.03 ± 0.0072 | 1.03 ± 0.0079 | 1.02 ± 0.0059 | 1.03 ± 0.0060 |
| **SfCc** | **Expected** | **6** | **2** | **1** | **3** |
|  | **AICc** | 1.85 ± 0.0555 | 1.30 ± 0.0317 | 1.07 ± 0.0124 | 2.12 ± 0.0757 |
|  | **KICc** | 1.66 ± 0.0515 | 1.22 ± 0.0289 | 1.04 ± 0.010 | 1.94 ± 0.070 |
|  | **BIC** | 1.49 ± 0.0476 | 1.13 ± 0.0248 | 1.02 ± 0.0039 | 1.71 ± 0.0637 |

M<sub>0</sub>: Random mating model. SfC: female competition and mate choice with independent parameters. SfCc: female competition and mate choice with compound parameters.

### Appendix D) Incomplete set-up: toy example

The proposed modelling framework requires at least two phenotypes by sex (mating tables of 2×2 dimensions or higher) for measuring sexual competition and mate choice effects. However it still can be applied if some sex, say females, have only one phenotype. In this case we just need to duplicate the row (see Fig. D1). Obviously, only male sexual selection can be measured.

**Fig. D1.**
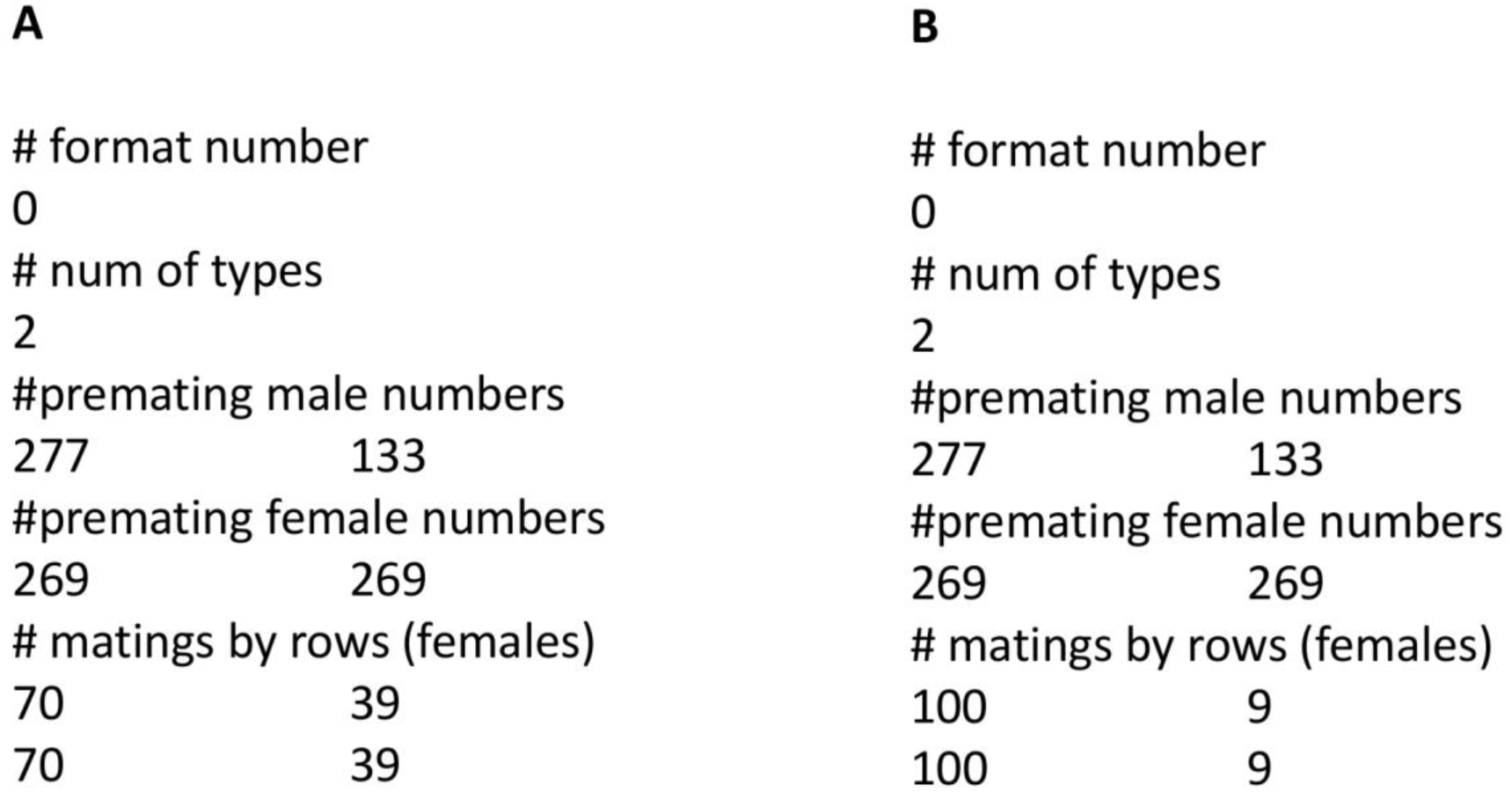
Examples of two toy models with only one type of female and two types of males. Note that the rows of the mating table are duplicated (same female type). A: Random mating B: Male sexual selection.

The examples in Fig. D1 correspond to a population with only one female but two male phenotypes (phenotype-1 and phenotype-2). There were sampled 269 females plus 277 males with phenotype-1 and 133 males with phenotype-2. In the first example (Fig. D1-A) there were 70 matings involving the male phenotype-1 and 39 with male phenotype-2. In the second example (Fig. D1-B) the matings were 100 with phenotype-1 and 9 with phenotype-2.

The analysis of the first case indicated that there was no significant deviation from random mating (*J*_PTI_ = 0.005, *P* = 0.78). The best model was the random mating model *M*_0_. As expected, the multi-model estimation of the mutual mating parameters was 1 for every parameter. The results were the same for the three information indices (AICc, KICc and BIC).

The analysis of the second case detected a deviation from random mating (*J*_PTI_ = 0.405, *P* < 10^-7^) due to male sexual selection (*J*_PS2_ = 0.405, *P* < 10^-7^) see (Carvajal-Rodríguez 2018b) for details of the *J* indices. The best model was male sexual selection with one parameter (Smale-1P). The male sexual selection component indicated five times higher mating propensity of male phenotype-1 with respect to phenotype-2.

## References

Aho, K., D. Derryberry, and T. Peterson. 2014. Model selection for ecologists: the worldviews of AIC and BIC. Ecology 95:631–636.

Aho, K., D. Derryberry, and T. Peterson. 2017. A graphical framework for model selection criteria and significance tests: refutation, confirmation and ecology. Methods in Ecology and Evolution 8:47–56.

Akaike, H. 1973. Information theory and an extension of the maximum likelihood principle. Pp. 267-281 *in* B. N. Petrov, and F. Csaki, eds. Second International Symposium on Information Theory, Budapest: Akademiai Kiado.

Anderson, D. R., K. P. Burnham, and G. C. White. 1994. AIC Model Selection in Overdispersed Capture-Recapture Data. Ecology 75:1780–1793.

Andersson, M. 1994. Sexual selection. Princeton University Press, Princeton, N.J.

Armstrong, D. M. 1977. Dispersal vs. Dispersion: Process vs. Pattern. Systematic Biology 26:210–211.

Arnold, S. J. and M. J. Wade. 1984. On the measurement of natural and sexual selection: applications. Evolution:720-734.

Barker, R. J. and W. A. Link. 2015. Truth, models, model sets, AIC, and multimodel inference: A Bayesian perspective. The Journal of Wildlife Management 79:730–738.

Brewer, M. J., A. Butler, and S. L. Cooksley. 2016. The relative performance of AIC, AICC and BIC in the presence of unobserved heterogeneity. Methods in Ecology and Evolution 7:679–692.

Burnham, K. P. and D. R. Anderson. 2002. Model selection and multimodel inference: a practical information-theoretic approach. Springer-Verlag, New York, NY.

Burnham, K. P., D. R. Anderson, and K. P. Huyvaert. 2011. AIC model selection and multimodel inference in behavioral ecology: some background, observations, and comparisons. Behavioral Ecology and Sociobiology 65:23–35.

Cade, B. S. 2015. Model averaging and muddled multimodel inferences. Ecology 96:2370–2382.

Carvajal-Rodríguez, A. 2018a. MateSim: Monte Carlo simulation for the generation of mating tables. Biosystems 171:26–30.

Carvajal-Rodríguez, A. 2018b. Non-random mating and information theory. Theoretical Population Biology 120:103–113.

Carvajal-Rodríguez, A. 2019. A generalization of the informational view of non-random mating: Models with variable population frequencies. Theoretical Population Biology 125:67–74.

Carvajal-Rodriguez, A. and E. Rolan-Alvarez. 2006. JMATING: a software for the analysis of sexual selection and sexual isolation effects from mating frequency data. BMC Evol Biol 6:40.

Casares, P., M. C. Carracedo, B. del Rio, R. Piñeiro, L. Garcia-Florez, and A. R. Barros. 1998. Disentangling the Effects of Mating Propensity and Mating Choice in Drosophila. Evolution 52:126–133.

Cavanaugh, J. E. 2004. Criteria for linear model selection based on Kullback’s symmetric divergence. Australian & New Zealand Journal of Statistics 46:257–274.

Claeskens, G. 2016. Statistical Model Choice. Annual Review of Statistics and Its Application 3:233–256.

Cruz, R., E. Rolán-Álvarez, and C. García. 2001. Sexual selection on phenotypic traits in a hybrid zone of Littorina saxatilis (Olivi). Journal of Evolutionary Biology 14:773–785.

Darwin, C. 1871. The descent of man, and selection in relation to sex. Murray.

Darwin, C. 1974. The Descent of Man, and Selection in Relation to Sex, London.

Dziak, J. J., D. L. Coffman, S. T. Lanza, R. Li, and L. S. Jermiin. 2019. Sensitivity and specificity of information criteria. bioRxiv:449751.

Edward, D. A. 2015. The description of mate choice. Behavioral Ecology 26:301–310.

Endler, J. A. 1986. Natural selection in the wild. Princeton University Press, Princeton, N.J.

Estévez, D., T. P. T. Ng, M. Fernández-Meirama, J. M. Voois, A. Carvajal-Rodríguez, G. A. Williams, J. Galindo, and E. Rolán-Alvarez. 2018. A novel method to estimate the spatial scale of mate choice in the wild. Behavioral Ecology and Sociobiology 72:195.

Fitze, P. S. and J.-F. L. Galliard. 2011. Inconsistency between Different Measures of Sexual Selection. The American Naturalist 178:256–268.

Futuyma, D. J. and M. Kirkpatrick. 2017. Evolution. Sunderland, Massachusetts U.S.A: Sinauer Associates, Inc. Publishers.

Galipaud, M., M. A. F. Gillingham, M. David, F. X. Dechaume-Moncharmont, and R. B. O’Hara. 2014. Ecologists overestimate the importance of predictor variables in model averaging: a plea for cautious interpretations. Methods in Ecology and Evolution 5:983–991.

Galipaud, M., M. A. F. Gillingham, and F.-X. Dechaume-Moncharmont. 2017. A farewell to the sum of Akaike weights: The benefits of alternative metrics for variable importance estimations in model selection. Methods in Ecology and Evolution 8:1668–1678.

Gao, F. and L. Han. 2012. Implementing the Nelder-Mead simplex algorithm with adaptive parameters. Computational Optimization and Applications 51:259–277.

Giam, X. and J. D. Olden. 2016. Quantifying variable importance in a multimodel inference framework. Methods in Ecology and Evolution 7:388–397.

Gimelfarb, A. 1988. Processes of Pair Formation Leading to Assortative Mating in Biological Populations: Encounter-Mating Model. The American Naturalist 131:865–884.

Grueber, C. E., S. Nakagawa, R. J. Laws, and I. G. Jamieson. 2011. Multimodel inference in ecology and evolution: challenges and solutions. Journal of Evolutionary Biology 24:699–711.

Hartl, D. L. and A. G. Clark. 1997. Principles of Population Genetics. Sinauer Associates, Inc., Sunderland, MA.

Jennions, M. D. and M. Petrie. 1997. Variation in mate choice and mating preferences: a review of causes and consequences. Biological Reviews 72:283.

Keerativibool, W. 2014. Unifying the Derivations of Kullback Information Criterion and Corrected Versions. Thailand Statistician 12:37–53.

Kim, H.-J., J. E. Cavanaugh, T. A. Dallas, and S. A. Foré. 2014. Model selection criteria for overdispersed data and their application to the characterization of a host-parasite relationship. Environmental and ecological statistics 21:329–350.

Kokko, H., H. Klug, and M. D. Jennions. 2012. Unifying cornerstones of sexual selection: operational sex ratio, Bateman gradient and the scope for competitive investment. Ecology Letters 15:1340–1351.

Lewontin, R., D. Kirk, and J. Crow. 1968. Selective mating, assortative mating, and inbreeding: definitions and implications. Eugen Q 15:141–143.

Link, W. A. and R. J. Barker. 2006. Model weights and the fundations of multimodel inference. Ecology 87:2626–2635.

Liu, W. and Y. Yang. 2011. Parametric or nonparametric? A parametricness index for model selection. The Annals of Statistics 39:2074–2102.

Mahler, D. L., M. G. Weber, C. E. Wagner, and T. Ingram. 2017. Pattern and Process in the Comparative Study of Convergent Evolution. The American Naturalist 190:S13–S28.

Ng, T. P. T., E. Rolán-Alvarez, S. S. Dahlén, M. S. Davies, D. Estévez, R. Stafford, and G. A. Williams. 2019. The causal relationship between sexual selection and sexual size dimorphism in marine gastropods. Animal Behaviour 148:53–62.

Parker, G. A. 2014. The sexual cascade and the rise of pre-ejaculatory (Darwinian) sexual selection, sex roles, and sexual conflict. Cold Spring Harbor perspectives in biology 6:a017509–a017509.

Parker, G. A. and T. Pizzari. 2015. Sexual Selection: The Logical Imperative. Pp. 119-163 *in* T. Hoquet, ed. Current Perspectives on Sexual Selection: What’s left after Darwin? Springer Netherlands, Dordrecht.

Press, W. H. 2002. Numerical recipes in C++: the art of scientific computing. CambridgeUniversity Press, Cambridge.

Prum, R. O. 2012. Aesthetic evolution by mate choice: Darwin’s really dangerous idea. Philosophical Transactions of the Royal Society of London B: Biological Sciences 367:2253–2265.

Richards, S. A. 2008. Dealing with overdispersed count data in applied ecology. Journal of Applied Ecology 45:218–227.

Rolan-Alvarez, E. 2007. Sympatric speciation as a by-product of ecological adaptation in the Galician Littorina saxatilis hybrid zone. Journal of Molluscan Studies 73:1–10.

Rolan-Alvarez, E., C. Austin, and E. G. Boulding. 2015a. The contribution of the genus Littorina to the field of evolutionary ecology. Oceanography and Marine Biology: an Annual Review 53:157–214.

Rolán-Alvarez, E. and A. Caballero. 2000. Estimating sexual selection and sexual isolation effects from mating frequencies. Evolution 54:30–36.

Rolan-Alvarez, E., A. Carvajal-Rodriguez, A. de Coo, B. Cortes, D. Estevez, M. Ferreira, R. Gonzalez, and A. D. Briscoe. 2015b. The scale-of-choice effect and how estimates of assortative mating in the wild can be biased due to heterogeneous samples. Evolution 69:1845–1857.

Rosenthal, G. G. 2017. Mate choice: the evolution of sexual decision making from microbes to humans. Princeton University Press.

Roughgarden, J., M. Oishi, and E. Akçay. 2006. Reproductive social behavior: cooperative games to replace sexual selection. Science 311:965–969.

Schwarz, G. 1978. Estimating the dimension of a model. The Annals of Statistics 6:461–464.

Shuker, D. M. 2010. Sexual selection: endless forms or tangled bank? Animal Behaviour 79:e11–e17.

Singer, S. and S. Singer. 2004. Efficient Implementation of the Nelder–Mead Search Algorithm. Applied Numerical Analysis & Computational Mathematics 1:524–534.

Symonds, M. R. E. and A. Moussalli. 2011. A brief guide to model selection, multimodel inference and model averaging in behavioural ecology using Akaike’s information criterion. Behavioral Ecology and Sociobiology 65:13–21.

Vrieze, S. I. 2012. Model selection and psychological theory: a discussion of the differences between the Akaike information criterion (AIC) and the Bayesian information criterion (BIC). Psychological methods 17:228.

Wacker, S. and T. Amundsen. 2014. Mate competition and resource competition are inter-related in sexual selection. Journal of evolutionary biology 27:466–477.

Wagenmakers, E.-J. 2007. A practical solution to the pervasive problems of p values. Psychonomic bulletin & review 14:779–804.

